# Identifying temporal and spatial patterns of variation from multi-modal data using MEFISTO

**DOI:** 10.1101/2020.11.03.366674

**Authors:** Britta Velten, Jana M. Braunger, Damien Arnol, Ricard Argelaguet, Oliver Stegle

## Abstract

Factor analysis is among the most-widely used methods for dimensionality reduction in genome biology, with applications from personalized health to single-cell studies. Existing implementations of factor analysis assume independence of the observed samples, an assumption that fails in emerging spatio-temporal profiling studies. Here, we present MEFISTO, a flexible and versatile toolbox for modelling high-dimensional data when spatial or temporal dependencies between the samples are known. MEFISTO maintains the established benefits of factor analysis for multi-modal data, but enables performing spatio-temporally informed dimensionality reduction, interpolation and separation of smooth from non-smooth patterns of variation. Moreover, MEFISTO can integrate multiple related datasets by simultaneously identifying and aligning the underlying patterns of variation in a data-driven manner. We demonstrate MEFISTO through applications to an evolutionary atlas of mammalian organ development, where the model reveals conserved and evolutionary diverged developmental programs. In applications to a longitudinal microbiome study in infants, birth mode and diet were highlighted as major causes for heterogeneity in the temporally-resolved microbiome over the first years of life. Finally, we demonstrate that the proposed framework can also be applied to spatially resolved transcriptomics.

## Introduction

Factor analysis is a first-line approach for the analysis of high-throughput sequencing data ^1–4^, and is increasingly applied in the context of multi-omics datasets ^5–8^. Given the popularity of factor analysis, this model class has undergone an evolution from conventional principal component analysis to sparse generalizations ^4^, including non-negativity constraints ^2,3,9^. Most recently, factor analysis has been extended to model structured datasets that consist of multiple data modalities or sample groups ^7,8^. At the same time, the complexity of multi-omics designs is constantly increasing, where in particular strategies for assaying multiple omics layers across temporal or spatial trajectories have gained relevance. However, existing factor analysis methods do not account for the resulting spatio-temporal dependencies between samples. Prominent domains where spatio-temporal designs are employed include developmental biology ^10^, longitudinal profiling in personalized medicine ^11^ or spatially resolved omics ^12^. Such designs and datasets pose new analytical challenges and questions, including (1) accounting for spatio-temporal dependencies across samples, which are no longer invariant to permutations, (2) dealing with imperfect alignment between samples from different data modalities and missing data, (3) identification of inter-individual heterogeneities of the underlying temporal and/or spatial functional modules and (4) distinguishing spatio-temporal variation from non-smooth patterns of variations. In addition, spatio-temporally informed dimensionality reduction can provide for more accurate and interpretable recovery of the underlying patterns, by leveraging known spatio-temporal dependencies rather than solely relying on feature correlations. To this end, we propose MEFISTO, a flexible and versatile method for addressing these challenges, while maintaining the benefits of previous factor analysis models for multi-modal data.

## Results

MEFISTO takes as input a dataset that contains measurements from one or more (possibly distinct) feature sets (e.g. different omics) - referred to as *views* in the following - as well as one or multiple sets of samples (e.g. from different experimental conditions, species or individuals) - referred to as *groups* in the following. In addition to this high-dimensional data, each sample is further characterized by a *continuous covariate* such as a one-dimensional temporal or two-dimensional spatial coordinate. MEFISTO factorizes the input data into *latent factors*, similar to conventional factor analysis, thereby recovering a joint embedding of the samples in a low-dimensional latent space. At the same time, the model yields a sparse and thus interpretable mapping between the latent factors and the observed features in terms of view-specific *weights*.

Critically, and unlike existing methods, MEFISTO incorporates the continuous covariate to account for spatio-temporal dependencies between samples, which allows for identifying both spatio-temporally smooth factors or non-smooth factors that are independent of the continuous covariate **(Figure 1A,B)**. Technically, MEFISTO combines factor analysis with the flexible non-parametric framework of Gaussian processes ^13^ to model spatio-temporal dependencies in the latent space, where each factor is governed by a continuous latent process to a degree depending on the factor’s smoothness (see **Supp. Methods**).

**Fig. 1.**
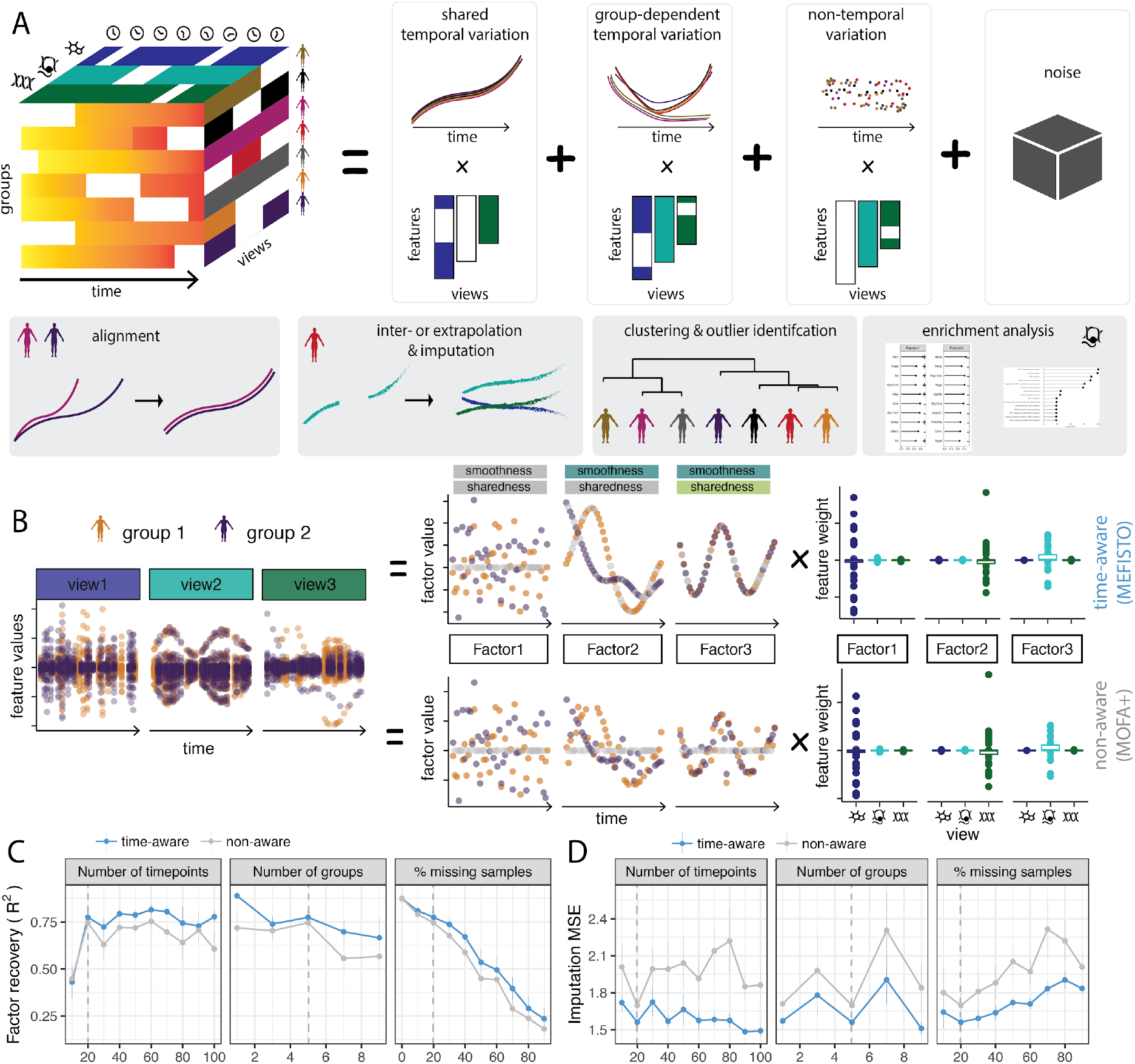
Method overview. **(A)** MEFISTO takes as input a possibly incomplete tensor-like high dimensional data set that can comprise multiple *views* (e.g. omics, tissues, genomic regions) measured in multiple sample *groups* (e.g. individuals, biological conditions, species) at multiple values of a covariate (e.g. time). Each element of this tensor is a high-dimensional feature vector of possibly different dimensions and non-matching features across views. The covariates can be misaligned across groups and arbitrary combinations of the dimensions can be missing. While accounting for the covariate, MEFISTO decomposes the high-dimensional input data into a set of smooth latent factors that capture temporal variation as well as latent factors that capture variation independent of the temporal axis. The latent factors can display arbitrary dependencies between groups including shared or group-specific factors. Sparse view-specific weights link a latent factor to individual views and features in the measurements. If covariates do not correspond between sample groups, MEFISTO infers a common scale by performing a simultaneous alignment and decomposition. Once learnt, the functional approach of MEFISTO enables novel downstream analyses, including interpolation or extrapolation and identification of relationships between sample groups or outliers per factor. **(B)** Illustration of a decomposition using MEFISTO compared to sparse factor analysis that is not aware of time (MOFA+) in a setting with a non-smooth factor (Factor 1), a smooth, non-shared factor (Factor 2) and a smooth, shared factor (Factor 3). **(C, D)** Comparison of MEFISTO (time-aware) to sparse factor analysis (non-aware, MOFA+) on simulated data in terms of recovery of the latent space (**C**) and mean squared error (MSE) of imputation for missing values (**D**) for varying number of time points, groups and level of missingness. The dashed line indicates the base parameters (see **Methods**) used for simulation.

For experimental designs with repeated spatio-temporal measurements, e.g. longitudinal studies involving multiple individuals, species or experimental conditions (termed *groups* in general), MEFISTO furthermore models and accounts for heterogeneity across these groups of samples, thereby inferring the extent to which spatio-temporal patterns are shared across groups (referred to as *sharedness*, **Figure 1B**). To cope with imperfect alignment across groups, MEFISTO comes with an integrated data-driven alignment step of the temporal covariate, e.g. aligning developmental stages between different species with unclear correspondences (see **Supp. Methods**).

To enable efficient inference in large datasets, MEFISTO leverages sparse Gaussian process approximations ^14^, as well as regular designs with a common spatio-temporal sampling across groups ^15^ (see **Supp. Methods)**. Once fitted, the model enables a broad range of downstream analyses (**Figure 1A**), including imputation as well as interpolation and extrapolation along the spatio-temporal axis, identification of molecular signatures underlying the latent factors using enrichment analysis as well as clustering and outlier identification on the level of samples, e.g. the measurement at a single time point, as well as groups of samples, e.g. an individual with distinct temporal trajectories.

### Validation using simulated data

To validate MEFISTO, we simulated time-resolved multi-modal data drawn from the generative model with multiple views and sample groups (**Methods**). We evaluated MEFISTO in terms of recovery of the latent factors, imputation of missing values in the high-dimensional input data, as well as recovery of the smoothness and sharedness of each factor. For comparison we also considered MOFA+ ^8^, a related multi-modal factor analysis method that however does not take the temporal covariate into account. Over a range of simulated settings, we observed improved recovery of the latent space and better imputation of missing data when accounting for the spatio-temporal dependencies (**Figure 1C,D**). At the same time, MEFISTO correctly determined the smoothness of the factors, allowing to distinguish temporal variation from non-temporal variation (**Supp. Fig. 1A**). In addition, the model correctly identified relationships of the groups, distinguishing group-specific and shared factors in a continuous manner (**Supp. Fig. 1B**). MEFISTO was robust to misaligned time points across the different sample groups, learning the correct alignment in a data-driven manner (**Supp. Fig. 2, 3**). Finally, we showed how the sparse Gaussian processes approximations employed by MEFISTO can improve its computational complexity, enabling applications to larger sample sizes, while maintaining accurate inference (**Supp. Fig. 4**).

**Figure 2:**
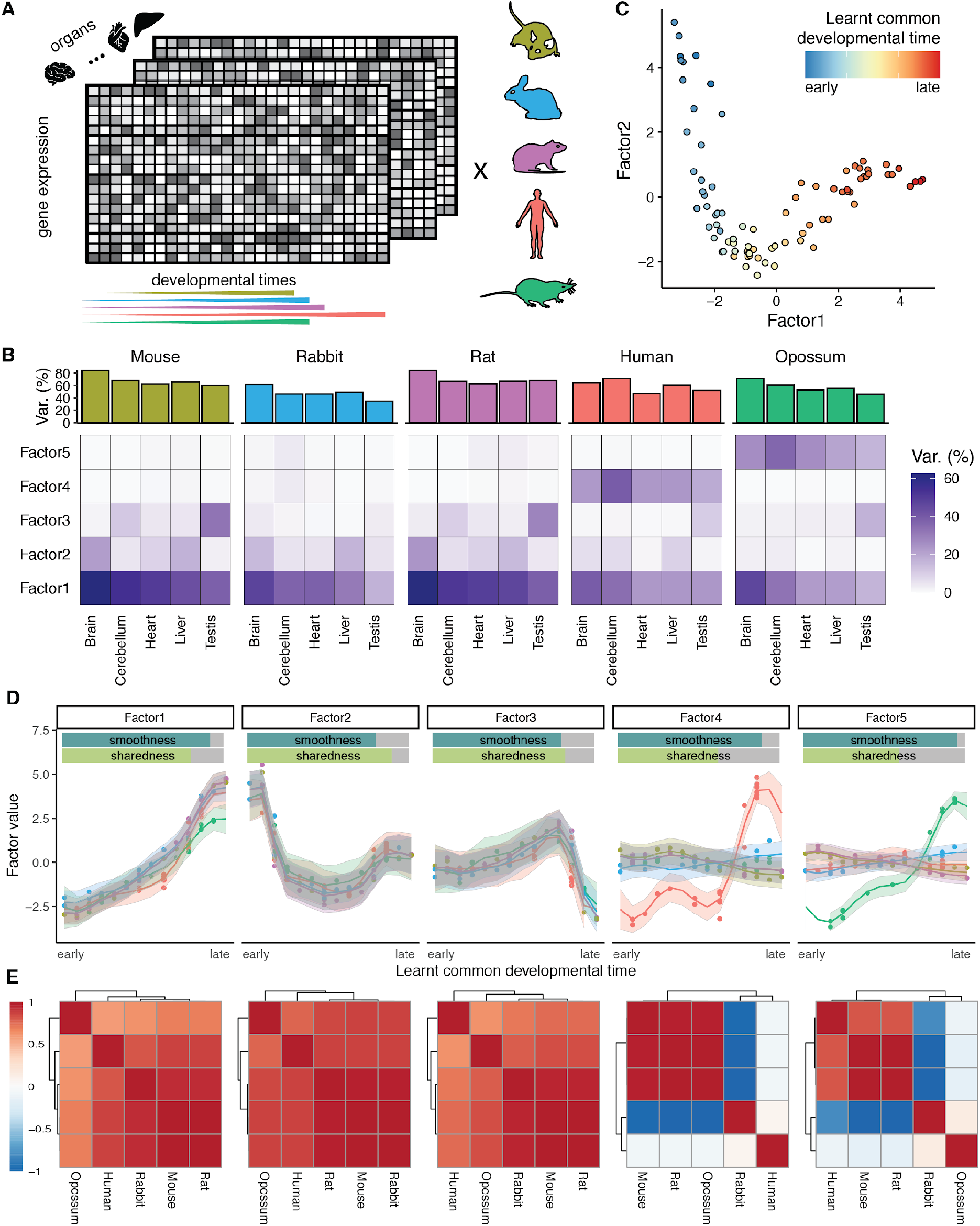
Application of MEFISTO to an evolutionary gene expression atlas across development. **(A)** Illustration of the input data and model setup, covering gene expression data from 5 species (groups) and 5 organs (views) across 14-23 developmental stages. **(B)** Variance explained in each species and organ per factor (bottom) and in total (top) **(C)** Scatter plot of first two latent factors with samples colored by the inferred common developmental time **(D)** Learnt latent factors are plotted against the inferred common developmental time. Points represent individual factor values, line and ribbons provide the mean and variance of the underlying latent process that generates the factor values. Bars on top indicate the smoothness along development and sharedness across species of the factor. **(E)** Learnt correlation structure of the species for the latent factor shown on top in **(D)**

**Figure 3.**
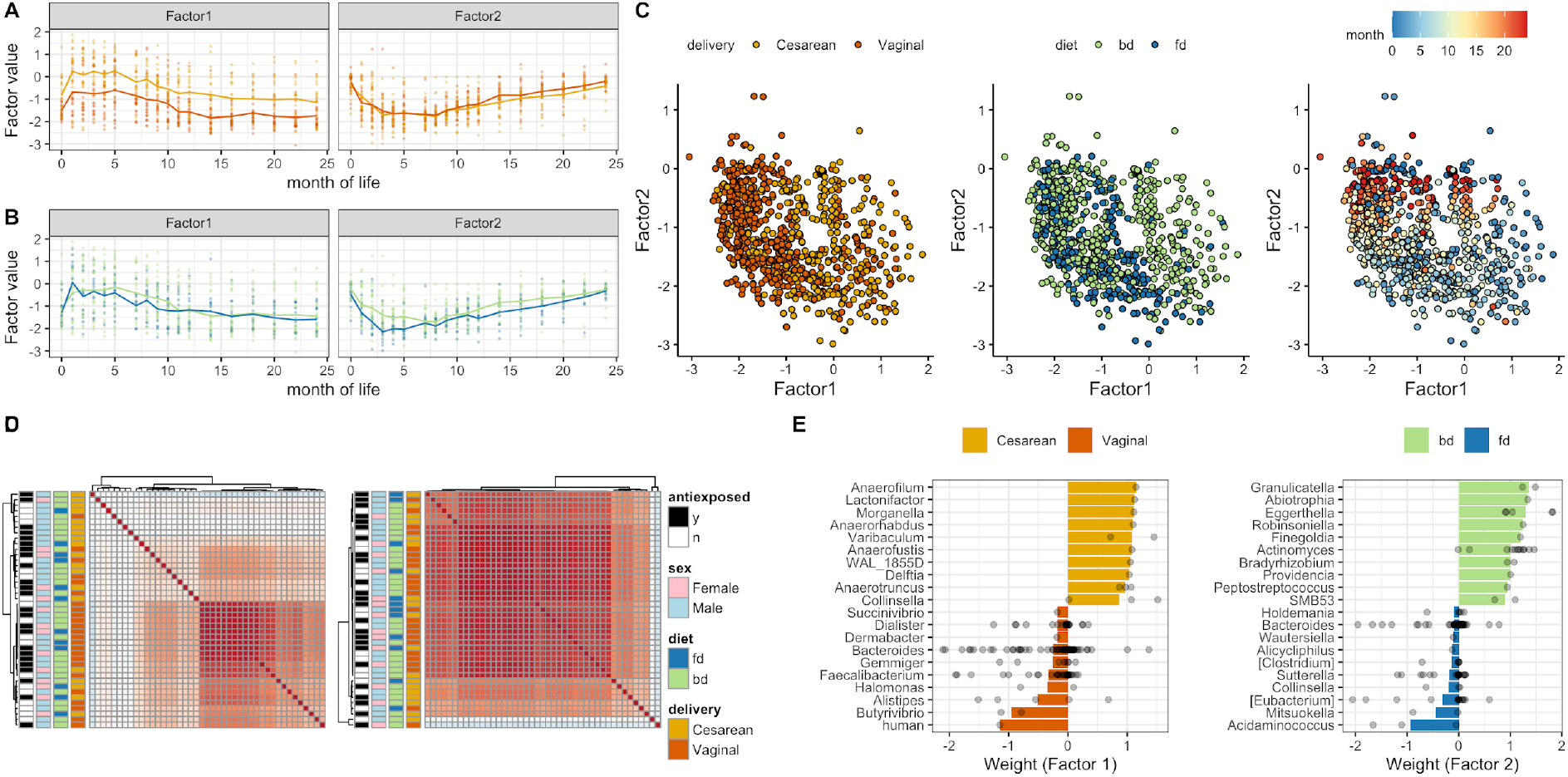
Application to a longitudinal microbiome study following children after birth. **(A, B)** Factor values (y-axis) across time (x-axis) colored by delivery mode **(A)** and predominant feeding mode **(B)**, termed diet; bd denotes breast milk-dominant, fd denotes formula-dominant. Dots represent inferred factor values per baby, the line shows the average across all samples in the respective category. **(C)** Scatterplot of samples on Factor 1 and 2 colored by delivery mode, diet and month of life **(D)** Inferred baby-baby correlation matrix for Factor 1 (left) and 2 (right) **(E)** Genus with the highest absolute mean weight for Factor 1 (left) and Factor 2 (right). Bars show the mean across the weights of all species in this genus, dots show the weights of individual species.

**Figure 4.**
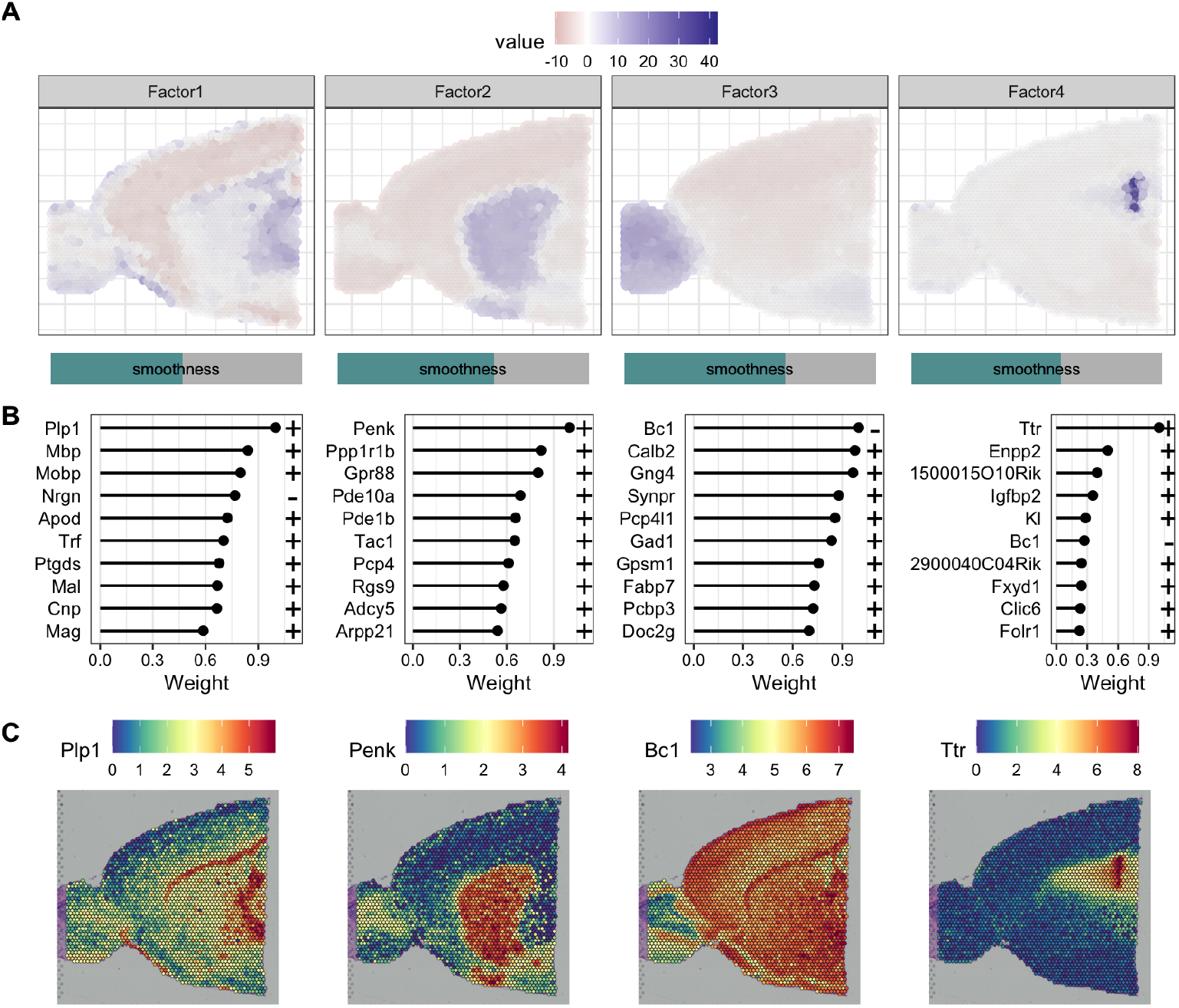
Application to spatial transcriptomics data. **(A)** Recovered factor values across space. Bars below indicate smoothness per factor. **(B)** Genes with highest absolute weight per factor **(C)** Normalized gene expression of genes with highest absolute weight per factor

### Application to an evolutionary atlas of mammalian organ development identifies conserved and diverged developmental programs

Next, we applied MEFISTO to a comprehensive evolutionary atlas of mammalian organ development ^10^ (**Figure 2A)**, comprising gene expression of 5 species (*groups*) profiled across 5 organs (*views*) along development, starting from early organogenesis to adulthood (14 - 23 time points per species). MEFISTO identified 5 robust latent factors (**Supp. Fig. 5**), which explained 35 to 85% of the transcriptome variation for different organs (**Figure 2B**). In addition to dealing with missing time points for several organs and species (**Supp. Fig. 6**), MEFISTO aligned the developmental time points of the samples (**Figure 2C, Supp. Fig. 7, 8**), thereby identifying meaningful correspondences of the developmental stages between species (**Supp. Fig. 9**). All five factors showed a high degree of smoothness (**Figure 2D**), which is consistent with most variation in this dataset being driven by developmental programs. However, the sharedness across species varied considerably between the factors (**Figure 2D**). The first three factors displayed similar temporal profiles across all species, indicating that they captured conserved development programs. Factor 1 explained variation in all organs (**Figure 2B**) and captured expression dynamics with a gradual change along development. To further characterize the underlying developmental programs and their molecular drivers, we investigated the weights of the factor. This revealed shared molecular signatures across organs that were linked to broad developmental processes and proliferation, e.g. cell cycle related pathways (**Supp. Fig. 10A**). Among the genes identified, there were key modulators of development such as *IGF2BP1, SOX11* or *KLF9* ^*16–18*^, which are ubiquitously expressed in all organs and display conserved expression dynamics across species (**Supp. Fig. 10B,C**). At the same time, the weights of Factor 1 also revealed conserved but organ-specific signatures that vary in line with the major functions of the respective organ, e.g. upregulation of *GFAP* along Factor 1 in brain tissues of all species (**Supp. Fig 11**)^19^. Factor 2 was also active in multiple organs (**Figure 2B**) and captured developmental programs with onset in intermediate development, for example as characterized by a transient upregulation of *HEMGN* during development in the liver along Factor 2 in all species (**Supp. Fig. 12**)^20^. Factor 3 captured conserved gene expression signatures specific to the development of testis with a sharp transition in gene expression with the onset of male meiosis (**Figure 2B, D, Supp. Fig. 13**). In addition to these shared factors, MEFISTO identified two factors that explained variation specific to some of the species (human and opossum, **Figure 2D**). Here, MEFISTO identified a clear clustering that separated either human (Factor 4) or opossum (Factor 5) from the other species, which is consistent with their larger evolutionary distances to the remaining species (**Figure 2E)**. Interestingly, these two affect gene expression programs in all organs **(Figure 2B, Supp. Fig. 14, 15)**. Based on the weights of these factors MEFISTO provides a direct means to identify genes for each organ that have undergone trajectory changes along evolution. To illustrate this, we compared the weights of these factors to previously identified genes with distinct developmental expression trajectories that evolved on the branch separating opossum or human from the other species and found a clear enrichment for high weights on the factors **(Supp. Fig. 16, 17)**. Finally, we considered this dataset to further assess the applicability of MEFISTO to settings with pronounced missingness: We masked data for random species - timepoint combinations in some or all of the organs, and observed an accurate imputation by MEFISTO and the ability to interpolate time points with completely missing data (**Supp. Fig 18**).

### Application to sparse longitudinal microbiome data

As a second use case, we applied MEFISTO to longitudinal samples of children’s microbiome after birth ^21,22^. As common in microbiome data and longitudinal studies, this dataset is extremely sparse with 97.1 - 99.8% of zero or missing values and up to 17 missing time points per child (out of 18, with an average of 4 time points missing). Nevertheless, MEFISTO identified distinct temporal trajectories depending on the birth mode (Factor 1, **Figure 3A**) and (to a lesser extent) the diet of the children (Factor 2, **Figure 3B**). Jointly, these two factors explained between 8 and 56% of variation in each child. While at the sample level clustering is mainly driven by time (**Figure 3C**), at the individual level Factor 1 shows a clustering depending on the delivery mode of the child (**Figure 3D**). Factor 2 does not show a clear clustering on the individual-level (**Figure 3D**). Microbial communities that are associated with Factor 1 reveal an enrichment of several bacteroid species in children with vaginal delivery as previously reported ^21,22^ (**Figure 3E**).

### Application to spatial transcriptomics

Last, we demonstrate the applicability of the method to spatial data by applying it to a spatial transcriptomics data set of anterior part of the mouse brain ^23^, where it identified major anatomical structures and its associated markers such as *Ttr* as a marker of the choroid plexus (**Figure 4**), without the need of single-cell reference data. In addition, MEFISTO provides an integrated measure of the smoothness of each pattern across space (**Figure 4A**). Making use of sparse inference, time and memory requirements could be greatly reduced compared to full inference (**Supp. Fig 19**).

## Conclusions

In summary, we here presented MEFISTO, a computational framework to open up multi-modal factor analysis models for applications to temporal or spatially-resolved data. We found that the ability to explicitly account for spatial or temporal dependencies is especially helpful in settings where data are sparse with many missing values in the high-dimensional measurements. Additionally, MEFISTO adds substantial value in cases where extra- or interpolation of temporal or spatial trajectories is required and/or when the temporal covariate and the associated measures are imperfectly aligned across data sets. We focused on an application of MEFISTO to temporal and longitudinal studies, such as the developmental time courses. These designs are rapidly gaining relevance both in basic biology and biomedicine. However, the model is also readily applicable to spatial domains and settings, as illustrated in the application to visium gene expression arrays. Future developments could focus on extensions to enable spatial alignment across datasets, as well as the deployment of specific noise models for example tailored for single-molecule data. In addition to time or space, other continuous covariates could be used to inform the factorization both for the factor values as well as their weights, e.g. using continuous clinical markers instead of time or known relationships in the feature space such as genomic positions in methylation or ATACseq data.

## Methods

### MEFISTO model

MEFISTO is a probabilistic model for factor analysis that accounts for continuous side information during inference of the latent space. To achieve this, MEFISTO combines multi-modal sparse factor analysis frameworks ^8^ with a functional view on the latent factors based on Gaussian processes and additionally provides alignment functionalities and an explicit model of inter-group heterogeneity. As input MEFISTO expects a collection of matrices, where each matrix ***Y***^***m,g***^ corresponds to a group *g* = 1, …, *G* and view *m* = 1, …, *M* with *N*_-_ samples in rows and *D*_*m*_ features in columns. Each sample is further characterized by a covariate 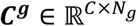 that represents for example temporal or spatial coordinates. The matrices are jointly decomposed as

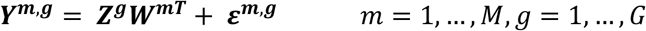

where 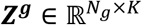 contains the *K* latent factors and 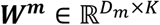 their weights. A feature- and view-wise sparsity prior is employed for ***W***^***m***^ as in previous multi-modal factor analyses models ^7,8^. Unlike existing factor models, however, the model additionally accounts for the covariate ***C***^***g***^. Each factor value 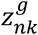 is modelled as realization of a Gaussian process

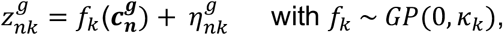

where the covariance function *K*_*k*_ models the relationship between groups as well as along the covariate, i.e.

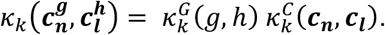

The first term in this covariance function captures the covariance of the discrete sample groups *g,h*, while the second term describes the covariance along values of the covariate, which provide a continuous characterization of each sample, e.g. its temporal or spatial location. We choose a low-rank covariance function for *K*^G^ and a squared exponential covariance function for *K*^C^, i.e.

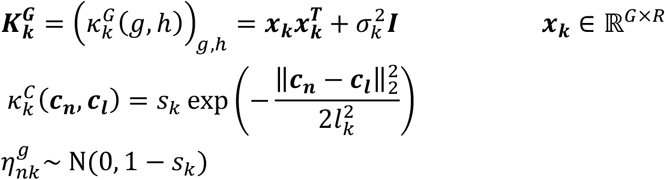

The hyperparameters ***x***_***k***_, *σ*_*k*_ *l*_*k*_, *S*_*k*_ determine the group-group covariance structure (***x***_***k***_, *σ*_*k*_) as well as the smoothness of the latent factors along the covariate (*l*_*k*_, *S*_*k*_). In particular, the scale parameter *S*_*k*_ determines the relative smooth versus non-smooth variation per factor, and the lengthscale parameter *l*_*k*_ the distance over which correlation decays along the covariate, e.g. in time or space. Details on the model specification, illustrations of the resulting covariance structures and a plate diagram are provided in **Supp. Methods, Section 2**.

### Inference

To infer the unobserved model components as well as the hyperparameters of the Gaussian process, MEFISTO makes use of variational inference combined with optimization of the evidence lower bound in terms of the hyperparameters of the Gaussian processes. Details on the inference are described in **Supp. Methods, Section 3**, where the specific updates of the inference algorithm are described. For large sample sizes, inference of the covariate kernel can be based on a subset of the original covariates chosen on a regular grid to reduce computational complexity (see **Supp. Methods, Section 4**). In addition, if the covariance matrix of the latent processes can be decomposed in terms of a Kronecker product, i.e. as ***K***^***G***^ ⨂ ***K***^***C***^, MEFISTO leverages this structure for accelerated inference based on spectral decomposition of the group- and covariate covariance (see **Supp. Methods, Section 3**).

### Alignment

If temporal correspondences between different groups are imperfect, a non-linear alignment between sample groups is learnt based on dynamic time warping ^24^ in the latent space. To reduce noise prior to the alignment, MEFISTO simultaneously decomposes the input data and aligns the covariate. This is implemented by interleaving the updates of the model components with an optimization step, where a warping curve is found that minimizes the distance of each group to a reference group in the current latent space. The alignment can be partial, i.e. have different end or start points between groups using an asymmetric step pattern in the time warping algorithm, or provide a global alignment using a symmetric step pattern in the time warping algorithm. Details are described in **Supp. Methods, Section 5**.

### Downstream analyses

Once the model is trained, the Gaussian process framework enables to interpolate or extrapolate the latent factors to unseen samples, groups or views as well as provide measures of uncertainty. Given a set of new covariate values ***c***^*^, MEFISTO makes predictions of the corresponding latent factor values ***z***^*^ based on the predictive distribution *p*(***z***^*^|***Y***) (see **Supp. Methods, Section 6**). Missing values of the considered features are then imputed from the model equation as in previous models ^7,8^. Furthermore, the hyperparameters of the model give insights into the smoothness of a factor (*S*_*k*_, between 0 (non-smooth) and 1 (smooth)) and the group relationships specific to a latent factor (***K***^***G***^) that can be used to cluster the groups or identify outliers. An overall sharedness score per factor is calculated by the mean absolute distance to the identity covariance matrix in the off-diagonal elements.

### Related methods

MEFISTO is related to previous matrix factorization and tensor decomposition methods, which however mostly ignore temporal information ^1–8^, use it only for pre-processing ^25^ or interpret it post-hoc ^22^. Those who incorporate such information do not allow multiple views (e.g. omics) ^26–28^ or are restricted to the same features in each view ^29^. In addition, sparsity constraints are not employed in these models which enhance interpretability and identifiability of the model. Besides linear methods, non-linear approaches have made use of continuous side-information, e.g. in the context of variational autoencoders ^30,31^ or recurrent neural networks ^32^. In particular, all of the above methods are incapable of handling non-aligned time courses across data sets (apart from Duncker & Sahani ^29^) and cannot capture heterogeneity across sample groups in the latent factors. For a detailed overview on related methods we refer to **Supp. Methods, Table 1**.

### Simulations

Data was simulated from the generative model varying the number of time points per group in a [0,1]-interval, noise levels, number of groups and fraction of missing values. Ten independent data sets were simulated for each setting from the generative model with three latent processes, having scale parameters of 1, 0.6, 0 and lengthscales of 0.2, 0.1, 0. For the first two (smooth and partially smooth) factors, one was randomly selected to be shared across all groups, while for the other factor a correlation matrix between groups of rank 1 was simulated randomly based on a uniformly distributed vector. MEFISTO was compared to MOFA+ ^8^ in terms of factor recovery given by the correlation of the inferred and simulated factor values as well as in terms of the mean squared error between imputed and ground-truth values for the masked values in the high-dimensional input data. Base settings for all non-varied parameters are 20 time points per group, five groups, four views with 500 features each and a noise variance of 1. 20% of randomly selected time points were masked per group and view, whereof 50% were missing in all views. To assess the alignment capabilities of the model, data was simulated with the same setup for 3 groups and covariates were transformed before training by a linear mapping (*h(t) = 0*.*4t +0*.*3*), a non-linear mapping (*h(t) = exp(t)*) and the identity in each group, respectively. These transformed covariates were passed to the model and the learnt alignment was compared to the ground-truth warping functions. To assess the scalability in the number of timepoints using sparse Gaussian processes, data was simulated from one group and with the same base parameters as above.

### Evo-devo data

Count data was obtained from Cardoso-Moreira et al ^10^, corrected for library size and normalised using variance stabilizing transformation provided by *DESeq2* ^*33*^. Genes were subsetted to orthologous genes as given in Cardoso-Moreira et al ^10^. Following the trajectory analysis of the original publication, we focused on 5 species, namely human, opossum, mouse, rat and rabbit, and 5 organs, namely brain, cerebellum, heart, liver and testis. In total, this resulted in a data set of 5 groups (species) and 5 views (organs) with 7,696 features each. The number of time points for each species varied between 14 and 23. As developmental correspondences were unclear we used a numeric ordering within each species ranging from 1 to the maximal number of time points in this species as input for MEFISTO and let the model infer the correspondences of time points between species. Stability analysis of the latent factors was performed by re-training the model on a down-sampled data set, where random selections of 1-5 time points were repeatedly masked in each organ-species combination. Gene set enrichment analysis was performed based on the reactome gene sets ^34^. To assess the imputation performance gene expression data in 2-20 randomly selected species - time combinations (out of a total of 82) were masked in 3, 4 or all organs and the model was retrained on this data as described above. The experiment was repeated ten times and the mean squared error was calculated on all masked values.

### Microbiome

Data was obtained from ‘Code Ocean’ capsule: https://doi.org/10.24433/CO.5938114.v1 and processed using a robust-centered log ratio and filtering steps as provided by Martino et al ^22^. This resulted in a total of 43 babies (*groups*) with up to 18 time points (months) and 3,236 features that were provided as input to MEFISTO using month of life as covariate. All zero values were treated as missing following previous work ^22^.

### Spatial transcriptomics

Data was obtained from the *SeuratData* R package as *stxBrain*.*anterior1*, normalized and subset to the 2,000 most variable features using *NormalizeData* and *FindVariableFeatures* functions provided by *Seurat* ^23^. Normalized expression values at all 2,696 spots were provided to MEFISTO with tissue coordinates as 2-dimensional covariate. For training of MEFISTO 1,000 inducing points were selected on a regular grid in space. For comparison a model with 500 inducing points and one with all spots was trained and compared in terms of their inferred factors.

### Data availability

The evodevo data was obtained from Cardoso-Moreira et al ^10^ and can be accessed from ArrayExpress with codes <u>E-MTAB-6782</u> (rabbit), <u>E-MTAB-6798</u> (mouse), <u>E-MTAB-6811</u> (rat), <u>E-MTAB-6814</u> (human) and <u>E-MTAB-6833</u> (opossum) (https://www.ebi.ac.uk/arrayexpress/). The microbiome data is based on Bokulich et al ^21^ and can be found on Qiita (http://qiita.microbio.me), processed data was obtained from the ‘Code Ocean’ capsule: https://doi.org/10.24433/CO.5938114.v1 provided by Martino et al ^22^. The spatial transcriptomics data set was obtained from the *SeuratData* package under the name *stxBrain*.*anterior1*.

### Code availability

MEFISTO is implemented as part of the MOFA framework ^7,8^ which is available at https://github.com/bioFAM/MOFA2. Installation instructions and tutorials can be found at https://biofam.github.io/MOFA2/MEFISTO. Code to reproduce all figures is available at https://github.com/bioFAM/MEFISTO_analyses. In addition, we provide vignettes on the main applications as part of the MEFISTO tutorials on https://biofam.github.io/MOFA2/MEFISTO.

## Acknowledgements

The authors thank Margarida Cardoso-Moreira for feedback on the evodevo application, Ilia Kats for helpful comments on the implementation and Haimasree Bhattacharya for testing the method on GPU. B.V. was funded by the BMBF (COMPLS project MOFA). D.A. and R.A. were funded by the EMBL PhD programme. O.S. was supported by core funding from EMBL, the German Cancer Research Center and funding from Chan Zuckerberg Initiative.

## Contributions

B.V., O.S., D.A. conceived the project.

B.V., D.A. and R.A. implemented the model.

B.V. analysed the data with contributions by J.B. on the evodevo data.

B.V. generated the figures.

B.V. and O.S. wrote the manuscript with input from all authors.

O.S. supervised the project.

## References

1. Stegle, O., Parts, L., Piipari, M., Winn, J. & Durbin, R. Using probabilistic estimation of expression residuals (PEER) to obtain increased power and interpretability of gene expression analyses. Nat. Protoc. 7, 500–507 (2012).

2. Gehring, J. S., Fischer, B., Lawrence, M. & Huber, W. SomaticSignatures: inferring mutational signatures from single-nucleotide variants. Bioinformatics 31, 3673–3675 (2015).

3. Alexandrov, L. B., Nik-Zainal, S., Wedge, D. C., Campbell, P. J. & Stratton, M. R. Deciphering signatures of mutational processes operative in human cancer. Cell Rep. 3, 246–259 (2013).

4. Witten, D. M., Tibshirani, R. & Hastie, T. A penalized matrix decomposition, with applications to sparse principal components and canonical correlation analysis. Biostatistics 10, 515–534 (2009).

5. Hore, V. et al. Tensor decomposition for multiple-tissue gene expression experiments. Nat. Genet. 48, 1094–1100 (2016).

6. Meng, C., Kuster, B., Culhane, A. C. & Gholami, A. M. A multivariate approach to the integration of multi-omics datasets. BMC Bioinformatics 15, 162 (2014).

7. Argelaguet, R., Velten, B., Arnol, D. & Dietrich, S. Multi-Omics Factor Analysis—a framework for unsupervised integration of multi-omics data sets. Mol. Syst. Biol. (2018).

8. Argelaguet, R. et al. MOFA+: a statistical framework for comprehensive integration of multi-modal single-cell data. Genome Biol. 21, 111 (2020).

9. Brunet, J.-P., Tamayo, P., Golub, T. R. & Mesirov, J. P. Metagenes and molecular pattern discovery using matrix factorization. Proc. Natl. Acad. Sci. U. S. A. 101, 4164–4169 (2004).

10. Cardoso-Moreira, M. et al. Gene expression across mammalian organ development. Nature 571, 505–509 (2019).

11. Schüssler-Fiorenza Rose, S. M. et al. A longitudinal big data approach for precision health. Nat. Med. 25, 792–804 (2019).

12. Ståhl, P. L. et al. Visualization and analysis of gene expression in tissue sections by spatial transcriptomics. Science 353, 78–82 (2016).

13. Rasmussen, C. E. & Williams, C. K. I. Gaussian Processes for Machine Learning. (University Press Group Limited, 2006).

14. Hensman, J., Fusi, N. & Lawrence, N. D. Gaussian Processes for Big Data. arXiv [cs.LG] (2013).

15. Rakitsch, B., Lippert, C., Borgwardt, K. & Stegle, O. It is all in the noise: Efficient multi-task Gaussian process inference with structured residuals. in Advances in Neural Information Processing Systems 26 (eds. Burges, C. J. C., Bottou, L., Welling, M., Ghahramani, Z. & Weinberger, K. Q.) 1466–1474 (Curran Associates, Inc., 2013).

16. Huang, X. et al. Insulin-like growth factor 2 mRNA-binding protein 1 (IGF2BP1) in cancer. J. Hematol. Oncol. 11, 88 (2018).

17. Bhattaram, P. et al. Organogenesis relies on SoxC transcription factors for the survival of neural and mesenchymal progenitors. Nat. Commun. 1, 9 (2010).

18. Zeng, Z., Velarde, M. C., Simmen, F. A. & Simmen, R. C. M. Delayed parturition and altered myometrial progesterone receptor isoform A expression in mice null for Krüppel-like factor 9. Biol. Reprod. 78, 1029–1037 (2008).

19. Landry, C. F., Ivy, G. O. & Brown, I. R. Developmental expression of glial fibrillary acidic protein mRNA in the rat brain analyzed by in situ hybridization. J. Neurosci. Res. 25, 194–203 (1990).

20. Yang, L. V., Nicholson, R. H., Kaplan, J., Galy, A. & Li, L. Hemogen is a novel nuclear factor specifically expressed in mouse hematopoietic development and its human homologue EDAG maps to chromosome 9q22, a region containing breakpoints of hematological neoplasms. Mechanisms of Development vol. 104 105–111 (2001).

21. Bokulich, N. A. et al. Antibiotics, birth mode, and diet shape microbiome maturation during early life. Sci. Transl. Med. 8, 343ra82 (2016).

22. Martino, C. et al. Context-aware dimensionality reduction deconvolutes gut microbial community dynamics. Nat. Biotechnol. (2020) doi:10.1038/s41587-020-0660-7.

23. Stuart, T. et al. Comprehensive Integration of Single-Cell Data. Cell 177, 1888–1902.e21 (2019).

24. Giorgino, T. & Others. Computing and visualizing dynamic time warping alignments in R: the dtw package. J. Stat. Softw. 31, 1–24 (2009).

25. Straube, J., Gorse, A.-D., PROOF Centre of Excellence Team, Huang, B. E. & Lê Cao, K.-A. A Linear Mixed Model Spline Framework for Analysing Time Course ‘Omics’ Data. PLoS One 10, e0134540 (2015).

26. Ramsay, J. & Silverman, B. W. Functional Data Analysis. (Springer Science & Business Media, 2013).

27. Yu, B. M. et al. Gaussian-process factor analysis for low-dimensional single-trial analysis of neural population activity. in Advances in Neural Information Processing Systems 21 (eds. Koller, D., Schuurmans, D., Bengio, Y. & Bottou, L.) 1881–1888 (Curran Associates, Inc., 2009).

28. Luttinen, J. & Ilin, A. Variational Gaussian-process factor analysis for modeling spatio-temporal data. in Advances in Neural Information Processing Systems 22 (eds. Bengio, Y., Schuurmans, D., Lafferty, J. D., Williams, C. K. I. & Culotta, A.) 1177–1185 (Curran Associates, Inc., 2009).

29. Duncker, L. & Sahani, M. Temporal alignment and latent Gaussian process factor inference in population spike trains. in Advances in Neural Information Processing Systems 31 (eds. Bengio, S. et al.) 10445–10455 (Curran Associates, Inc., 2018).

30. Casale, F. P., Dalca, A., Saglietti, L., Listgarten, J. & Fusi, N. Gaussian Process Prior Variational Autoencoders. in Advances in Neural Information Processing Systems 31 (eds. Bengio, S.et al.) 10369–10380 (Curran Associates, Inc., 2018).

31. Fortuin, V., Baranchuk, D., Raetsch, G. & Mandt, S. GP-VAE: Deep Probabilistic Time Series Imputation. in (eds. Chiappa, S. & Calandra, R.) vol. 108 1651–1661 (PMLR, 2020).

32. Qiu, L., Chinchilli, V. M. & Lin, L. Deep Latent Variable Model for Longitudinal Group Factor Analysis. arXiv [stat.ML] (2020).

33. Love, M. I., Huber, W. & Anders, S. Moderated estimation of fold change and dispersion for RNA-seq data with DESeq2. Genome Biol. 15, 550 (2014).

34. Croft, D. et al. The Reactome pathway knowledgebase. Nucleic Acids Res. 42, D472–7 (2014).

