## Supplementary Figures for "Identifying temporal and spatial patterns of variation from multi-modal data using MEFISTO"

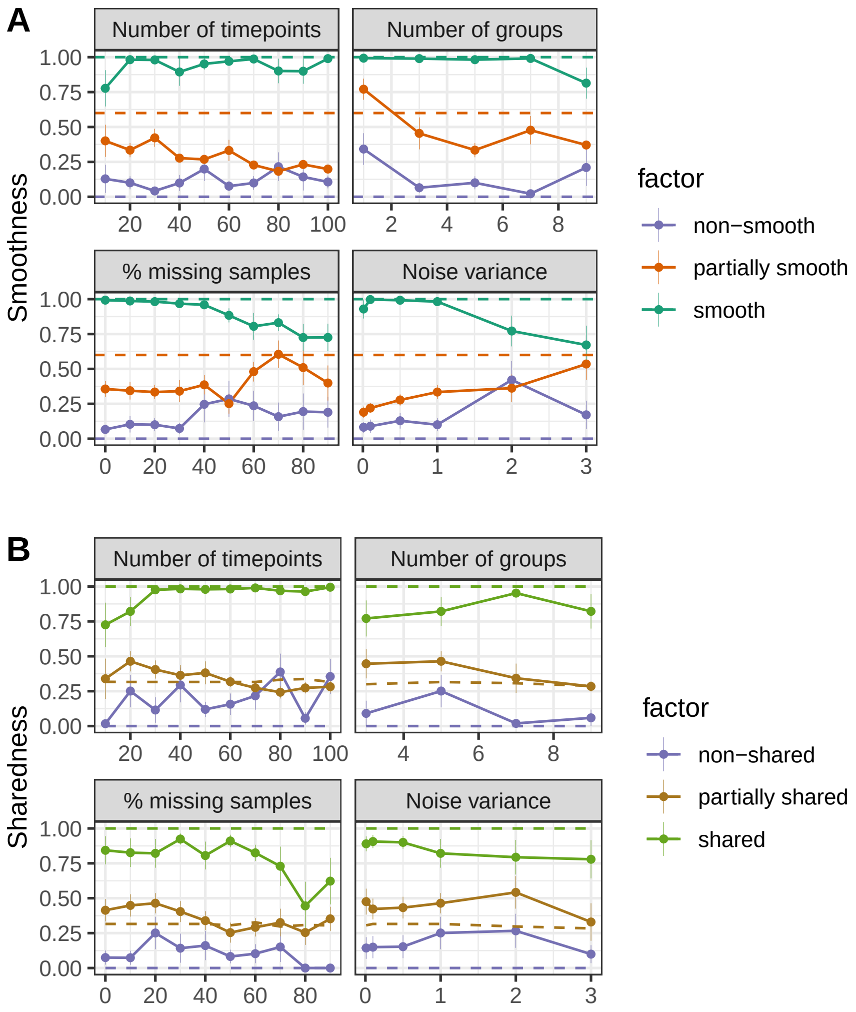


**Supp. Fig. 1: Inference of factor smoothness and sharedness on simulated data** For varying number of time points per group, noise levels, number of groups and fractions of missing values ten independent data sets were simulated from the generative model with three factors in each setting. The three factors were simulated to have different smoothness along time and sharedness across groups as indicated by the dashed lines (see **Methods**). Solid lines and dots show the smoothness score (A) and the sharedness score (B) inferred by MEFISTO per factor. Base settings for all non-varied parameters are 20 time points per group, 5 groups, 4 views with 500 features each, a noise variance of 1 and 20% of randomly selected time points missing per group and view, whereof 50% are missing in all views.

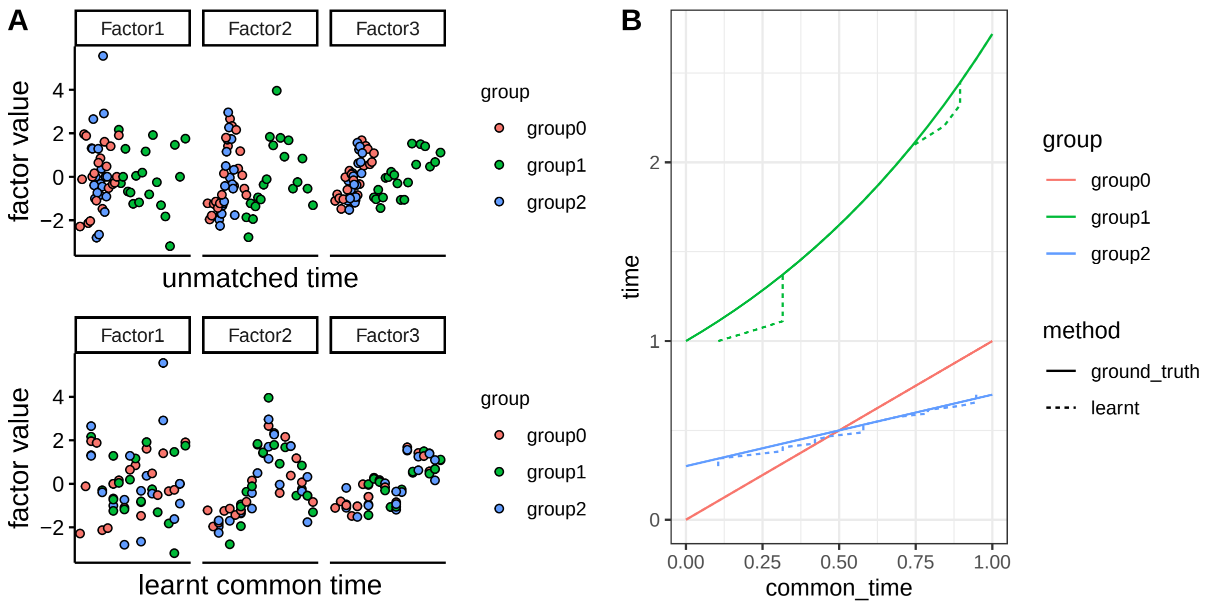


**Supp. Fig. 2: Illustration of alignment.** Illustration of MEFISTO's alignment on one example data set with 3 groups and 20 time points per group. Data was simulated with base parameters as described in Supp. Figure 1 and one non-smooth and two shared, smooth factors. (A) shows the learnt factor values (y-axis) against the observed time (x-axis, top) and the learnt common time (x-axis, bottom). (B) shows the learnt warping function per group (dashed line) compared to the ground truth (solid line).

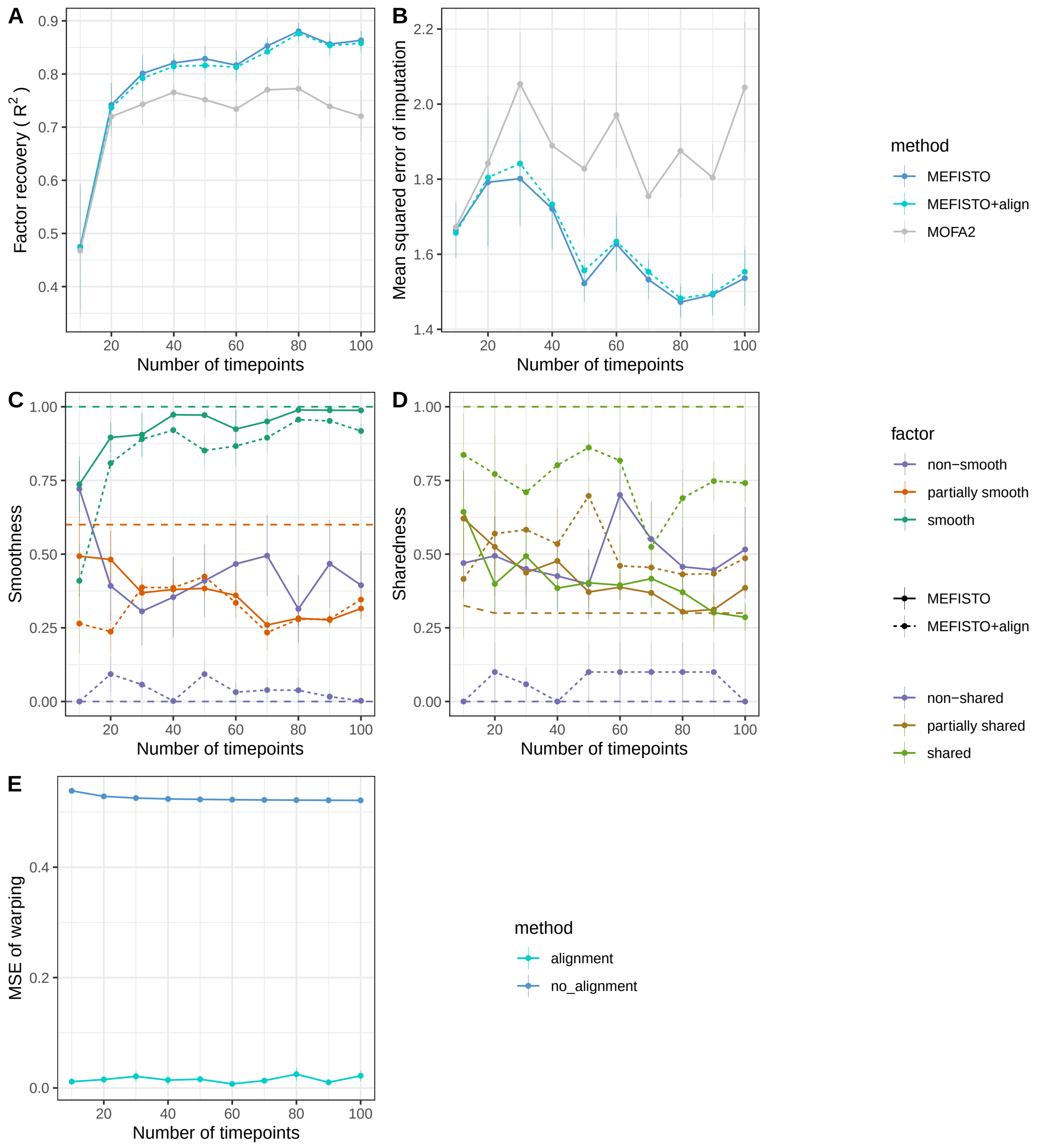


**Supp. Fig. 3: Validation of alignment on simulated data.** For three groups and varying number of time points per group (x-axis) ten independent data sets were simulated from an unobserved common time as in Supp. Figure 1. Afterwards, observed times per group were obtained from group-specific transformations of the unobserved common time via linear ($f(t) = 0.4\,t + 0.3$), exponential ($f(t) = \exp(t)$) or identity ($f(t) = t$) transformation. MOFA+, MEFISTO without alignment and MEFISTO with alignment were compared in terms of (A) overall factor recovery, (B) imputation mean squared error, (C) smoothness and (D) sharedness inference per factor as well as (E) quality of the learnt warping function.

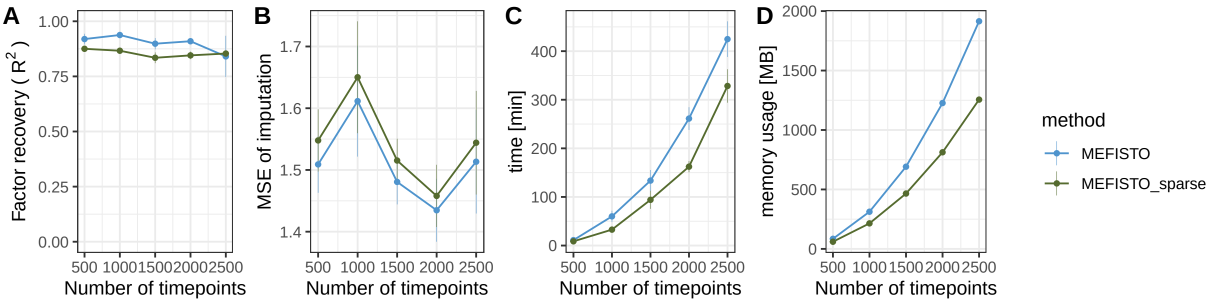


**Supp. Fig. 4: Validation of sparse Gaussian processes on simulated data** Factor recovery (A), mean squared error of imputation (B), time (C) and memory usage (D) compared for MEFISTO and a sparse version of MEFISTO using 75 % of total sample size as inducing points for varying number of time points (x-axis) and a single group. Remaining parameters for simulations are as described in Supp. Figure 1.

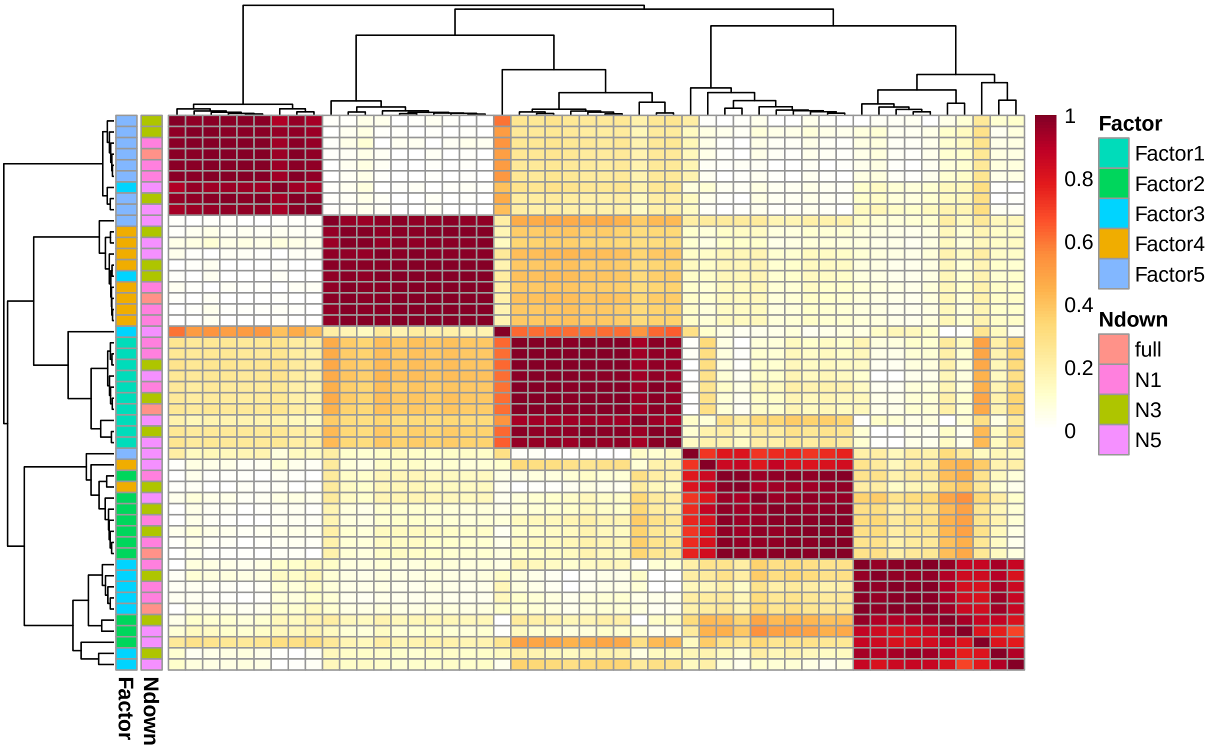


**Supp. Fig. 5: Factor stability in the evodevo application** Pearson correlation of factor values for models trained on the full data and on data where 1-5 time points for every organ and species have been downsampled. The five blocks on the diagonal indicate that each factor is robustly found in all models.

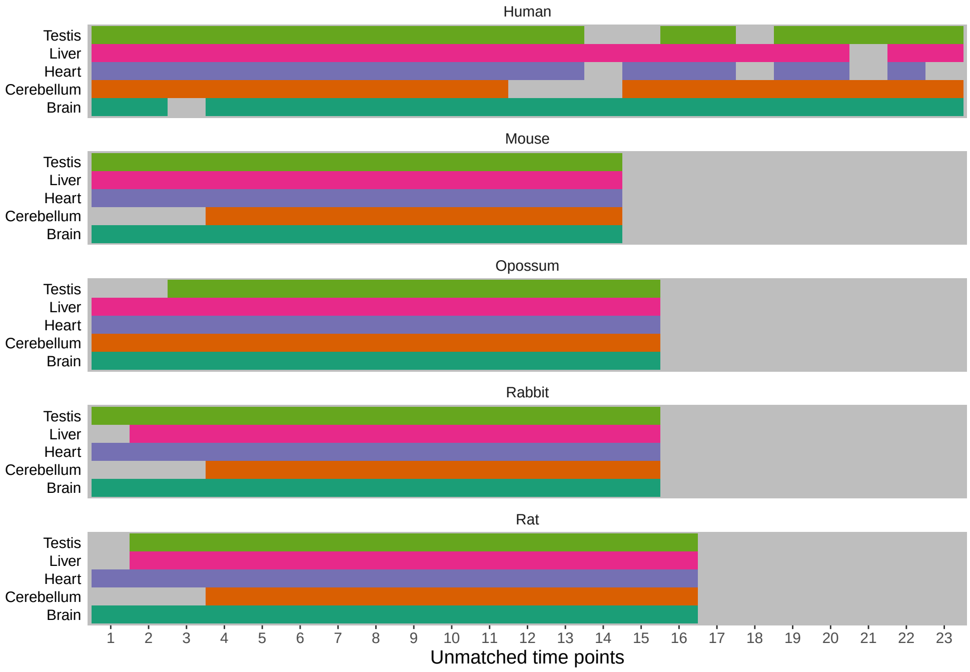


**Supp. Fig. 6: Data overview for the evodevo application** For each of the five species, gene expression data for 7,696 orthologous genes in five organs (y-axis, colours) were provided as views to MEFISTO. The x-axis shows the samples per species (ordered by developmental stage with numeric time points used as covariate in the model). Grey areas indicate missing samples. The developmental stages corresponding to the time points are shown in Supp. Figure 9.

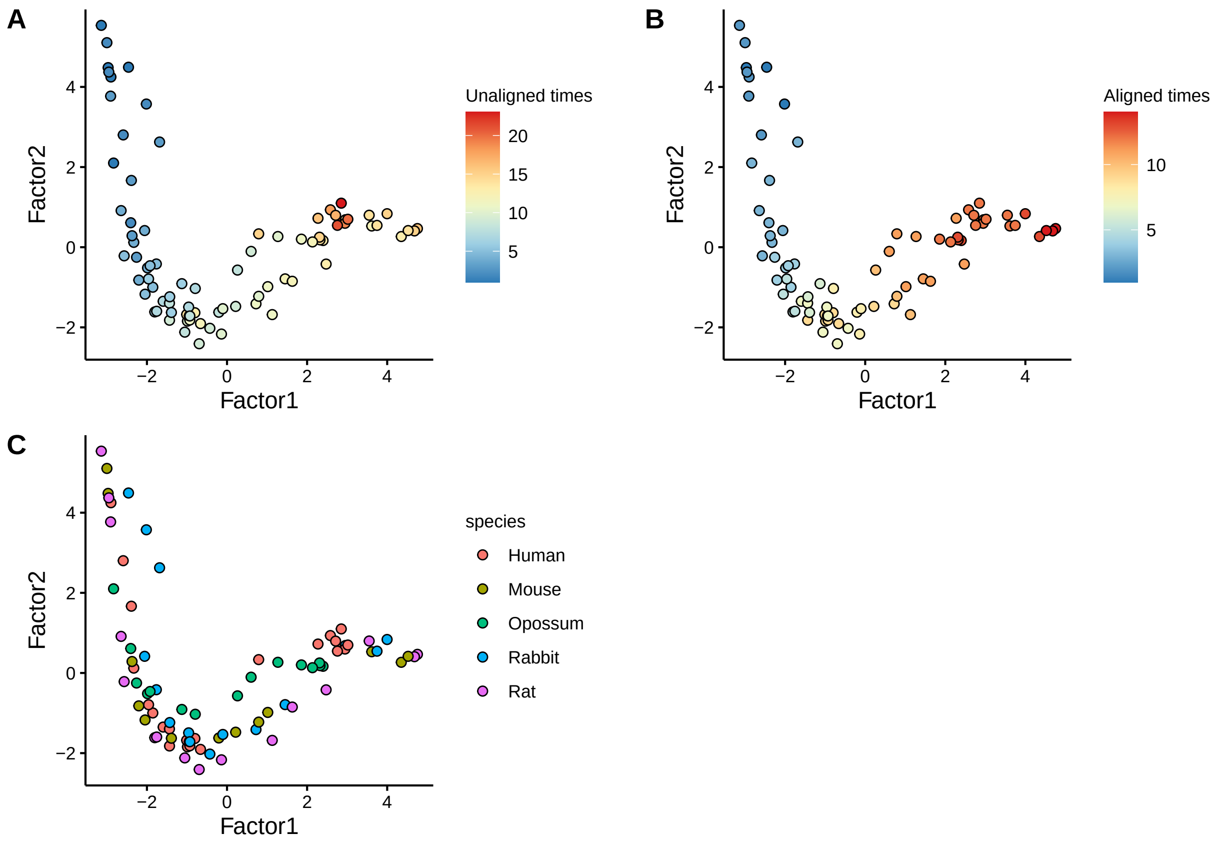


**Supp. Fig. 7: Latent embedding given by the first two factors** Each dot represent the factor values of the first (x-axis) and second (y-axis) factor for one species- timepoint combination. Dots are coloured by unaligned times (A), aligned times (B) and species (C).

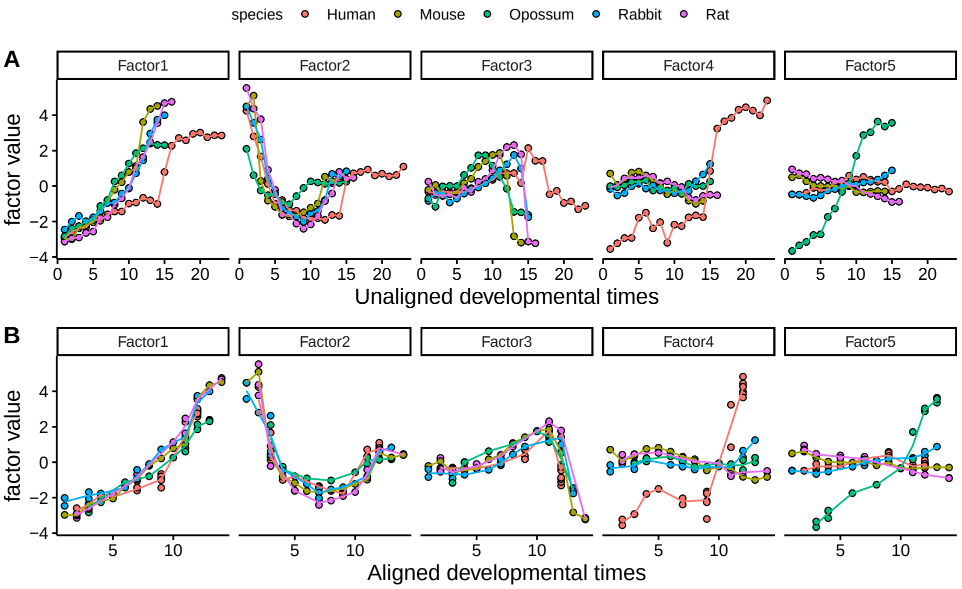


**Supp. Fig. 8: Factor values along time with and without alignment** (A) shows the factor values against the developmental stages without alignment across species, (B) shows the factor values against the developmental stages with alignment across species.

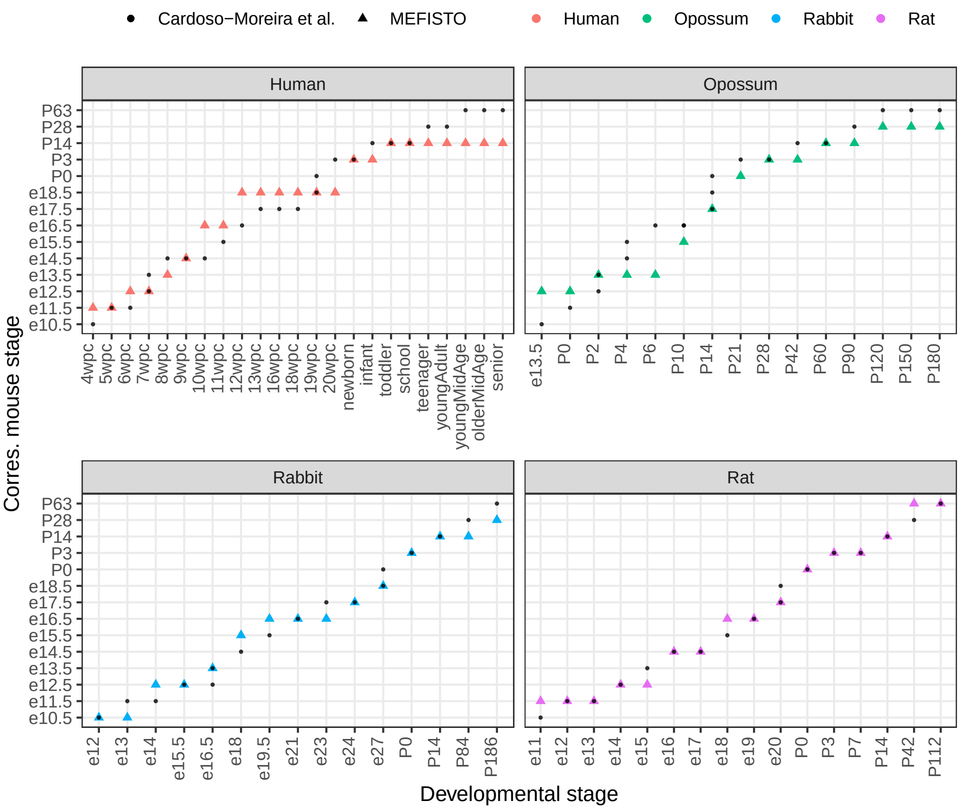


**Supp. Fig. 9: Alignment of developmental stages across species** For the developmental stages (x-axis) of each species (panels and colours) the corresponding mouse stage is shown on the y-axis. Triangles show the learnt correspondences, small black dots the developmental time correspondences according to Cardoso-Moreira et al.

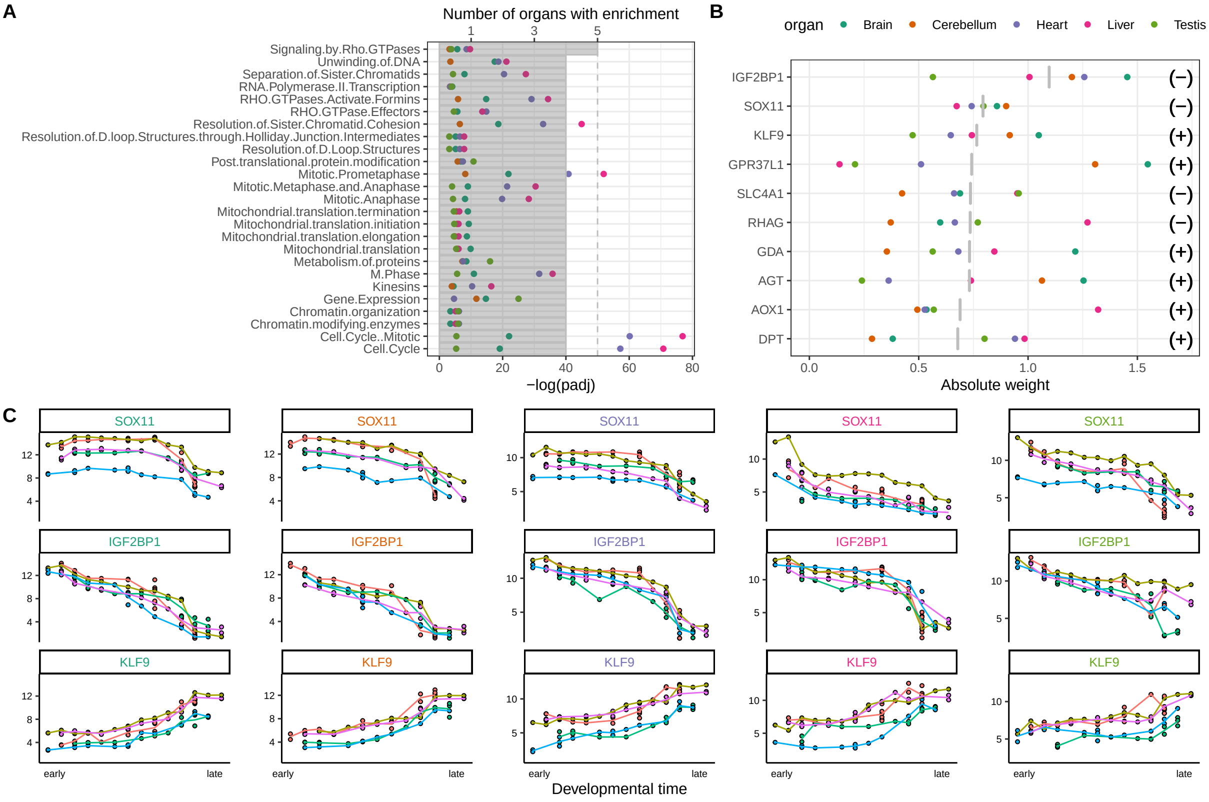


**Supp. Fig. 10: Pan-organ developmental programs on Factor 1 in the evodevo application** (A) Gene sets at a false discovery rate of 5% that are enriched in the weights of Factor 1 in at least 4 organs. Dots are coloured by organ and indicate the significance of a gene set (negative adjusted logarithmic p-value on the x-axis). Grey bars indicate the number of organs with enrichment. (B) Top 10 genes (y-axis) with highest absolute mean weight across organs. Dots are coloured by organ and indicate the absolute weight per organ, grey bar show the mean across organs. Signs on the right show the sign of the weights. (C) Gene expression along inferred developmental time in all organs (columns) for the top 3 genes of panel (B).

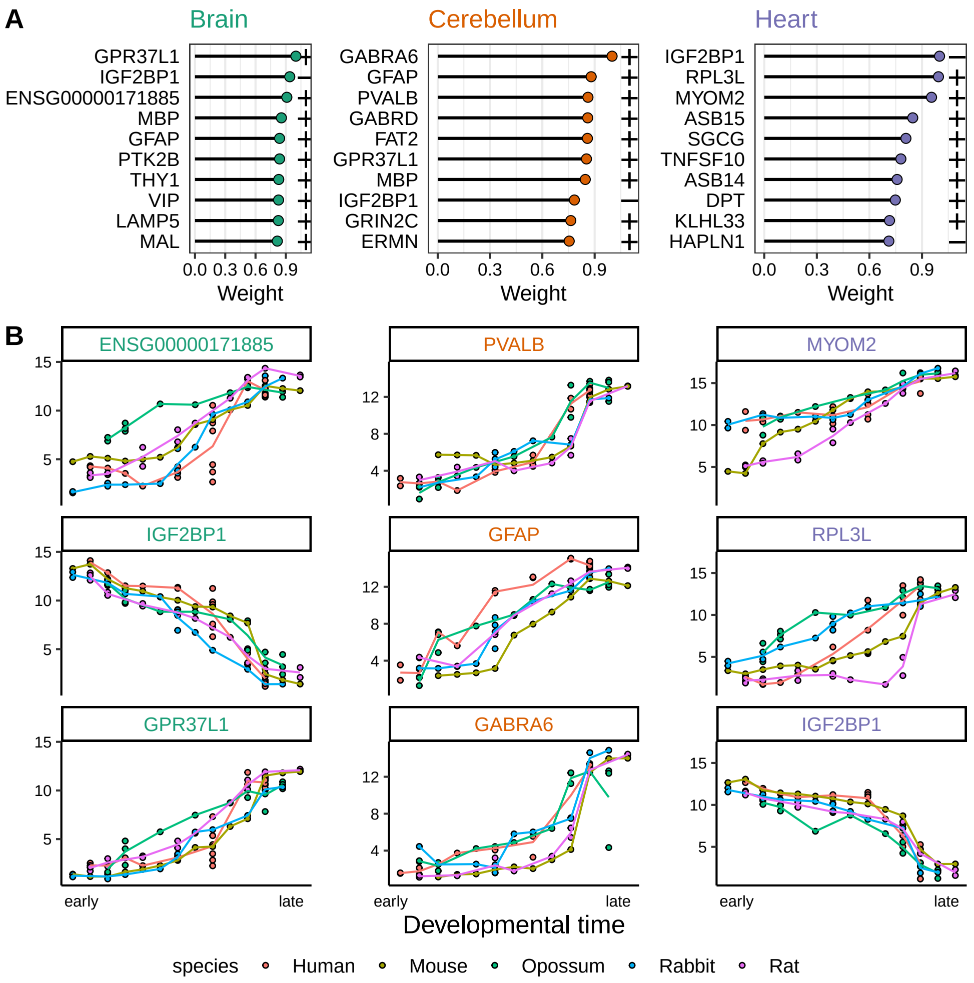


**Supp. Fig. 11: Organ-wise weights of Factor 1 in the evodevo application** (A) Genes with highest absolute weight (x-axis) for the three organs with highest variance explained by Factor 1. Symbols on the right in each panel indicate the sign of the weight. (B) Gene expression trajectories along inferred developmental time for the top 3 genes of the corresponding panel in (A).

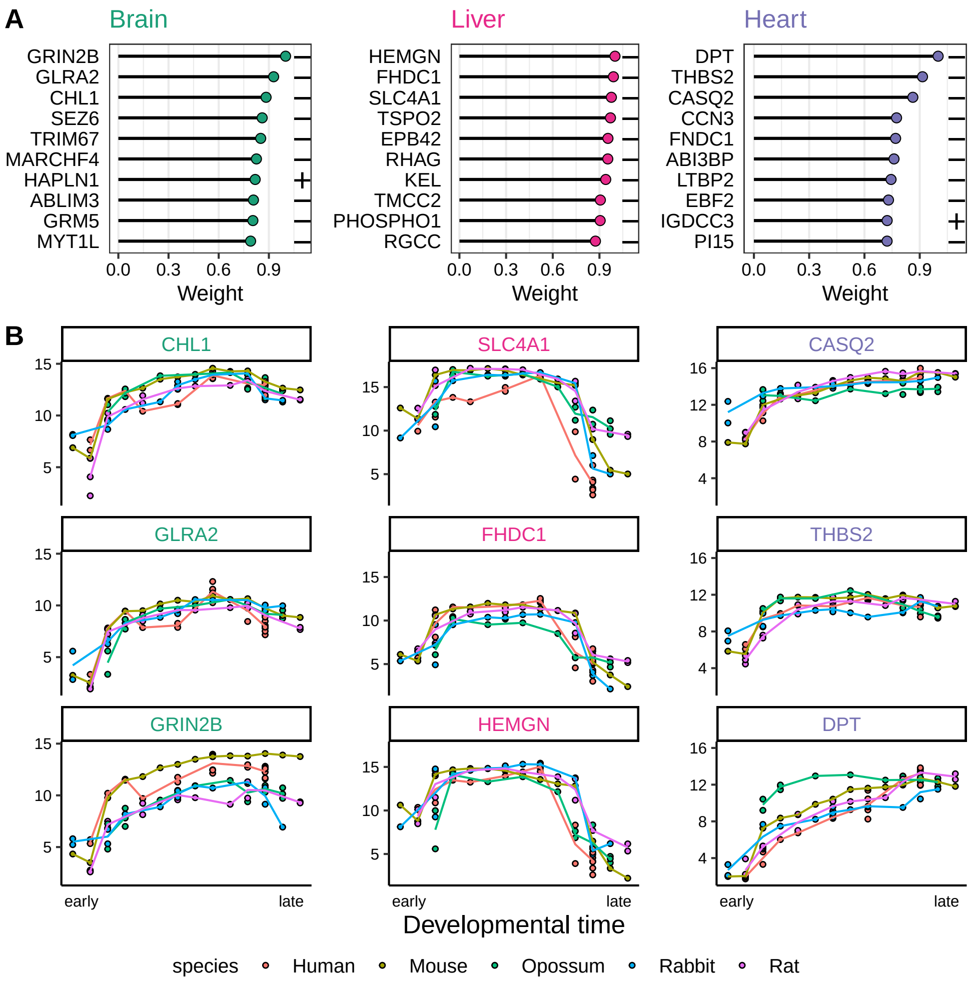


**Supp. Fig. 12: Organ-wise weights of Factor 2 in the evodevo application** (A) Genes with highest absolute weight (x-axis) for the three organs with highest variance explained by Factor 2. Symbols on the right in each panel indicate the sign of the weight. (B) Gene expression trajectories along inferred developmental time for the top 3 genes of the corresponding panel in (A).

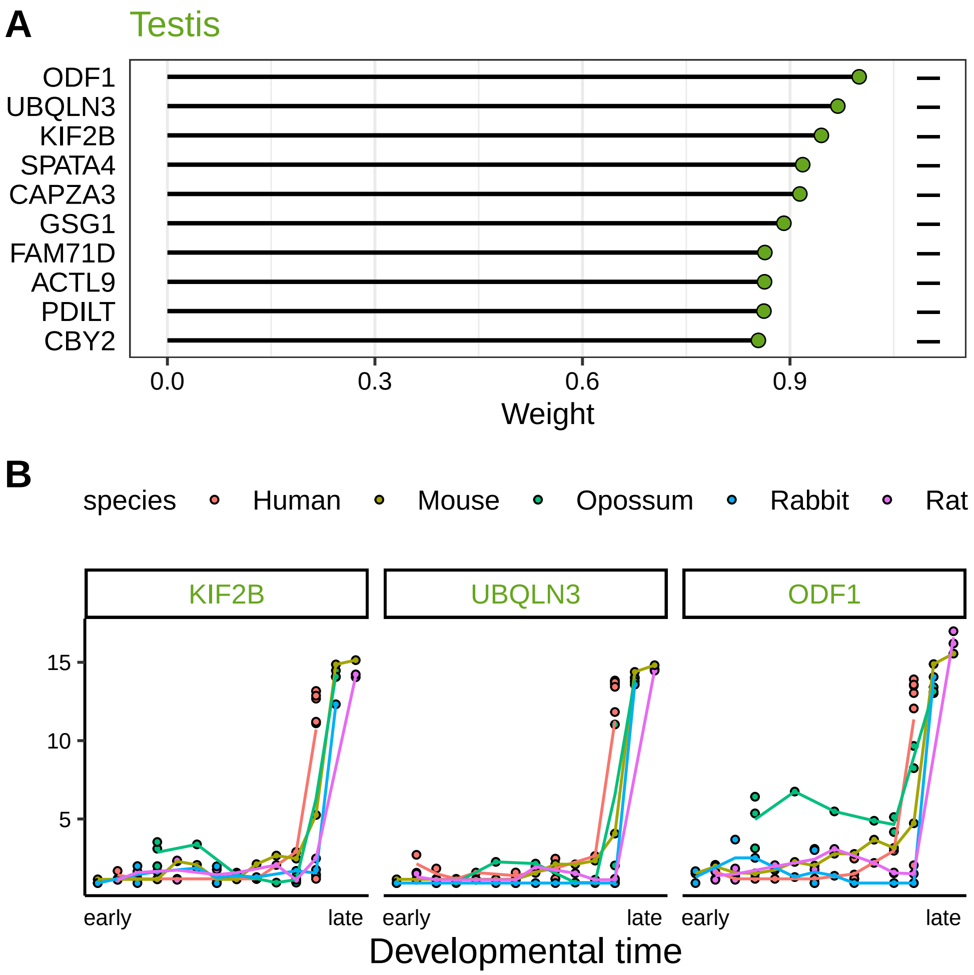


**Supp. Fig. 13: Testis weights of Factor 3 in the evodevo application** (A) Genes with highest absolute weight (x-axis) in Testis on Factor 3. Symbols on the right indicate the sign of the weight. (B) Gene expression trajectories along inferred developmental time for the top 3 genes in (A).

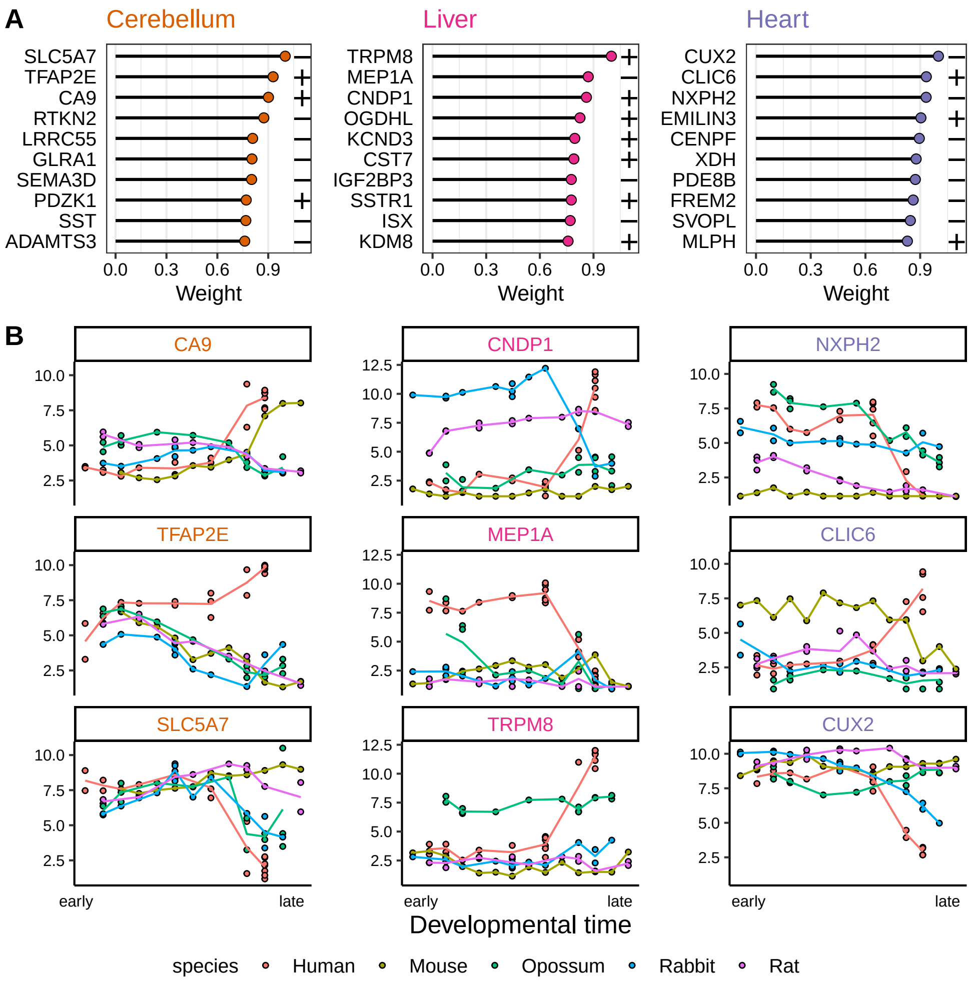


**Supp. Fig. 14: Organ-wise weights of Factor 4 in the evodevo application** (A) Genes with highest absolute weight (x-axis) for the three organs with highest variance explained by Factor 4. Symbols on the right in each panel indicate the sign of the weight. (B) Gene expression trajectories along inferred developmental time for the top 3 genes of the corresponding panel in (A).

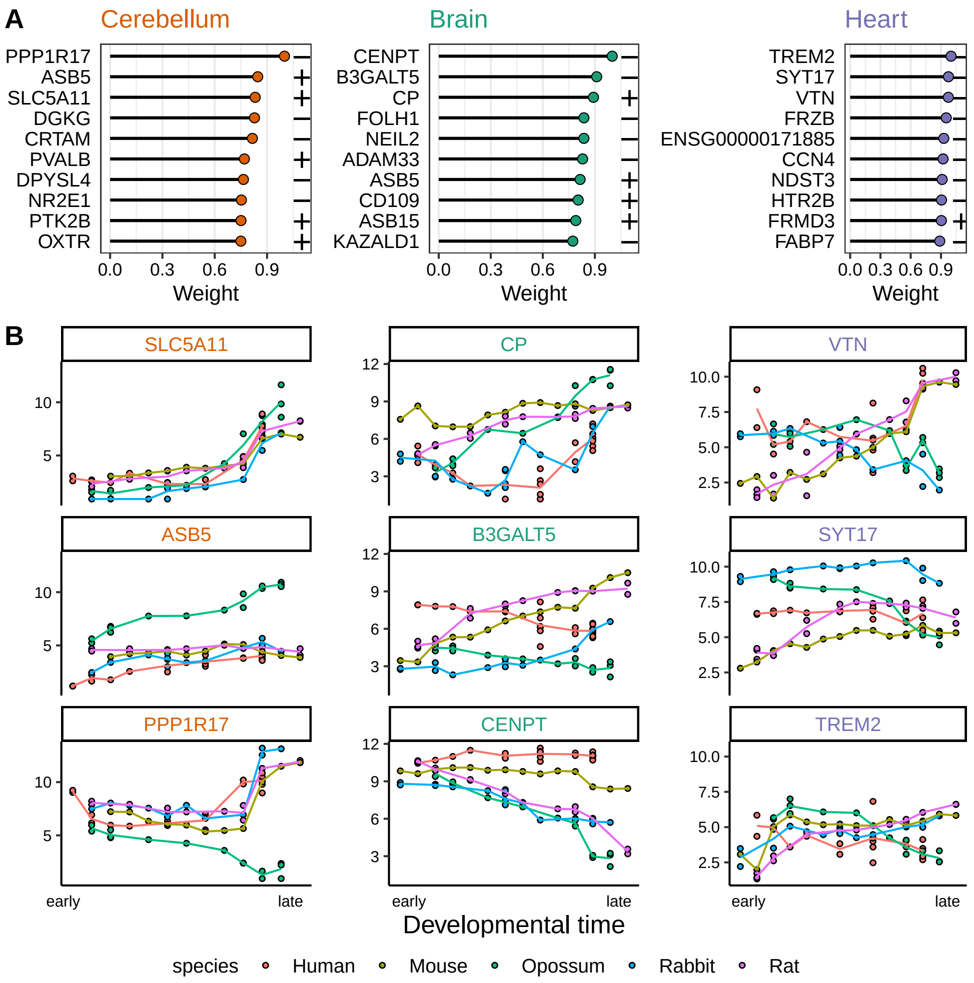


**Supp. Fig. 15: Organ-wise weights of Factor 5 in the evodevo application** (A) Genes with highest absolute weight (x-axis) for the three organs with highest variance explained by Factor 5. Symbols on the right in each panel indicate the sign of the weight. (B) Gene expression trajectories along inferred developmental time for the top 3 genes of the corresponding panel in (A).

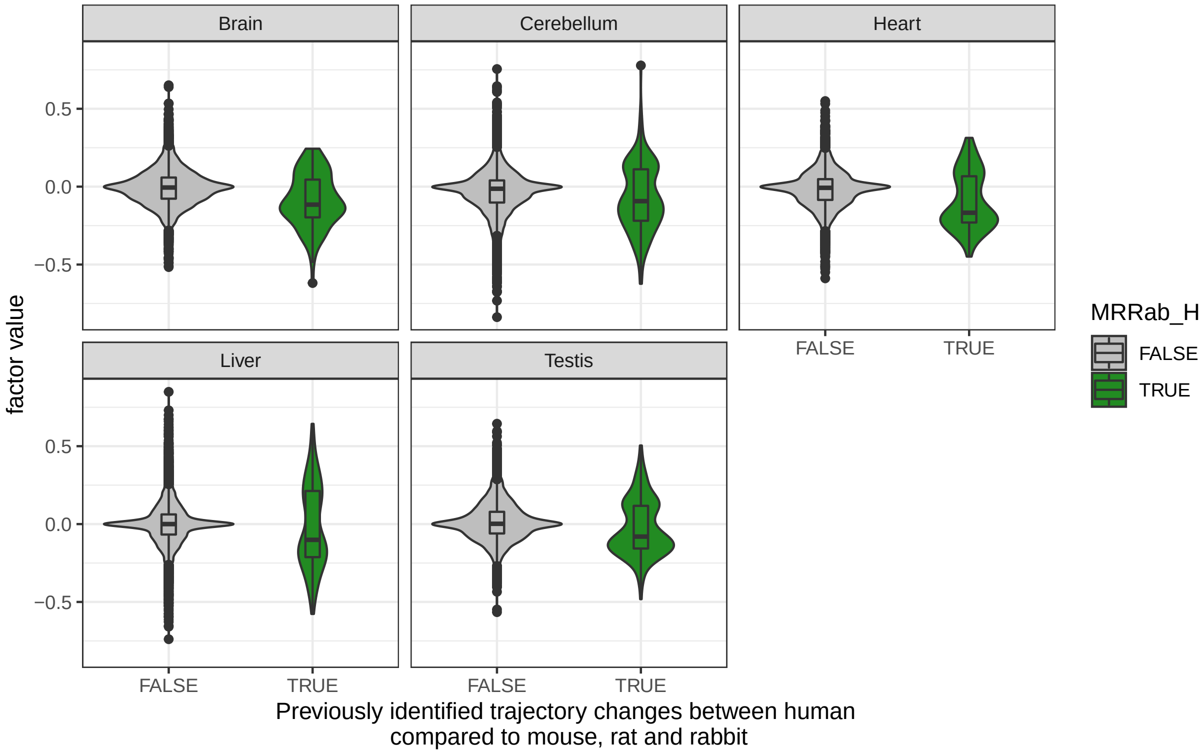


**Supp. Fig. 16: Weights of Factor 4 in the evodevo application split by classification in Cardoso-Moreira et al.** Shown are violin plots of the weights in the model for each organ (panels) separated by whether they have previously been identified as having changed developmental trajectories for human compared to rodents or rabbit (x-axis).

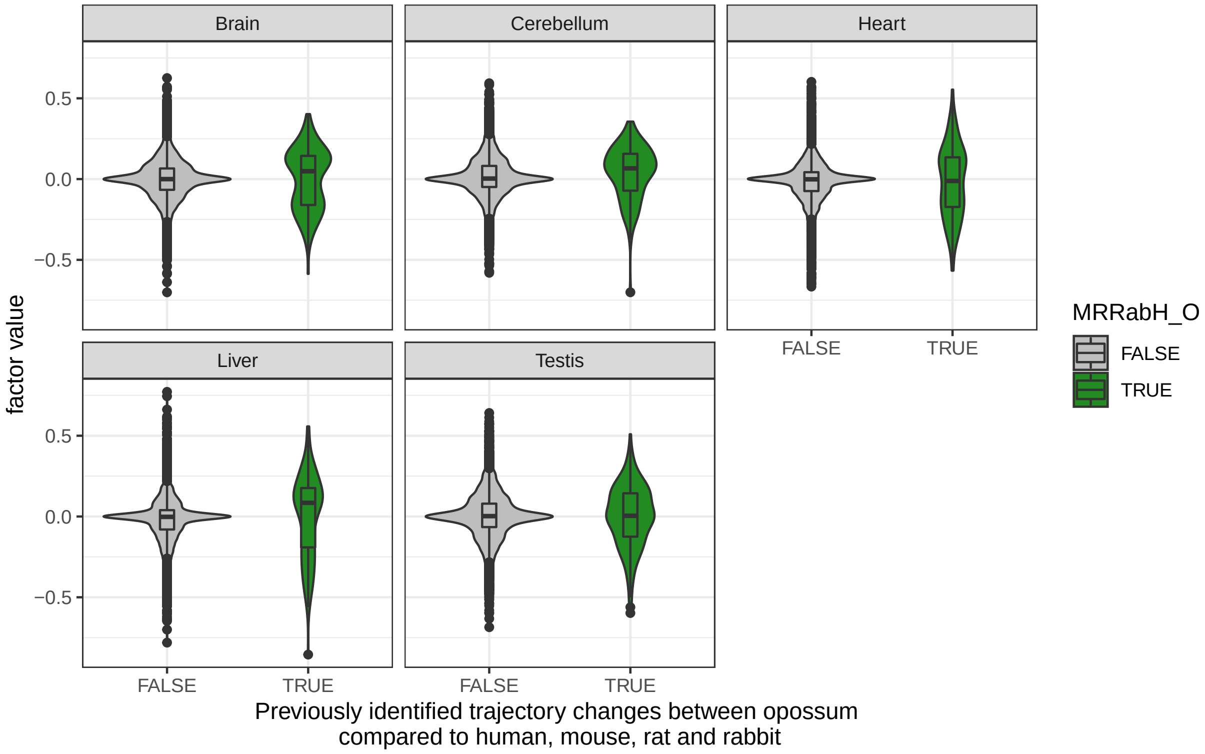


**Supp. Fig. 17: Weights of Factor 5 in the evodevo application split by classification in Cardoso-Moreira et al.** Shown are violin plots of the weights in the model for each organ (panels) separated by whether they have previously been identified as having changed developmental trajectories for opossum compared to the other mammals (x-axis).

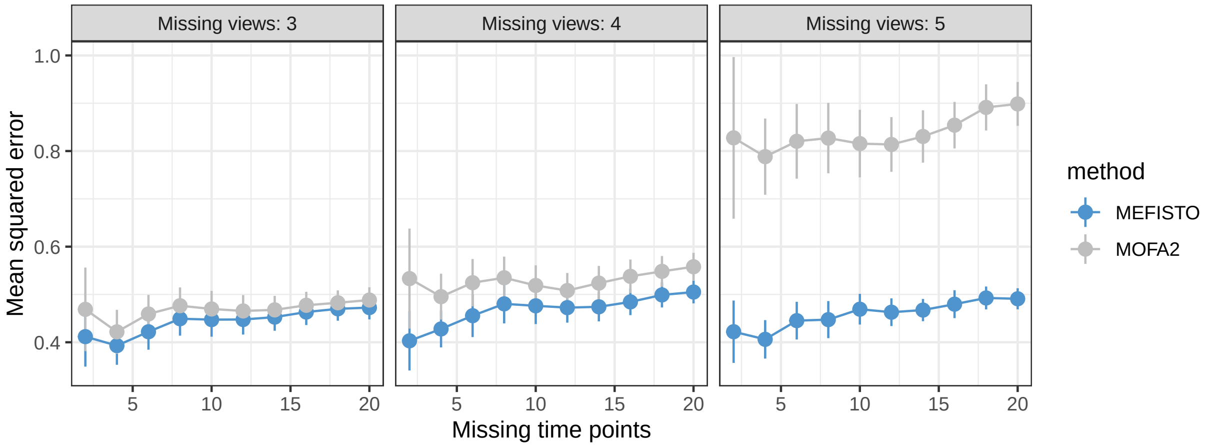


**Supp. Fig. 18: Interpolation experiment on the evodevo data** MOFA+ and MEFISTO were trained on the evodevo data after masking the expression data of all genes for a varying number of time points (x-axis, out of 82 available time points, 14-23 per species) in 3, 4 or all organs. The y-axis shows the mean squared error of imputation on all masked values.

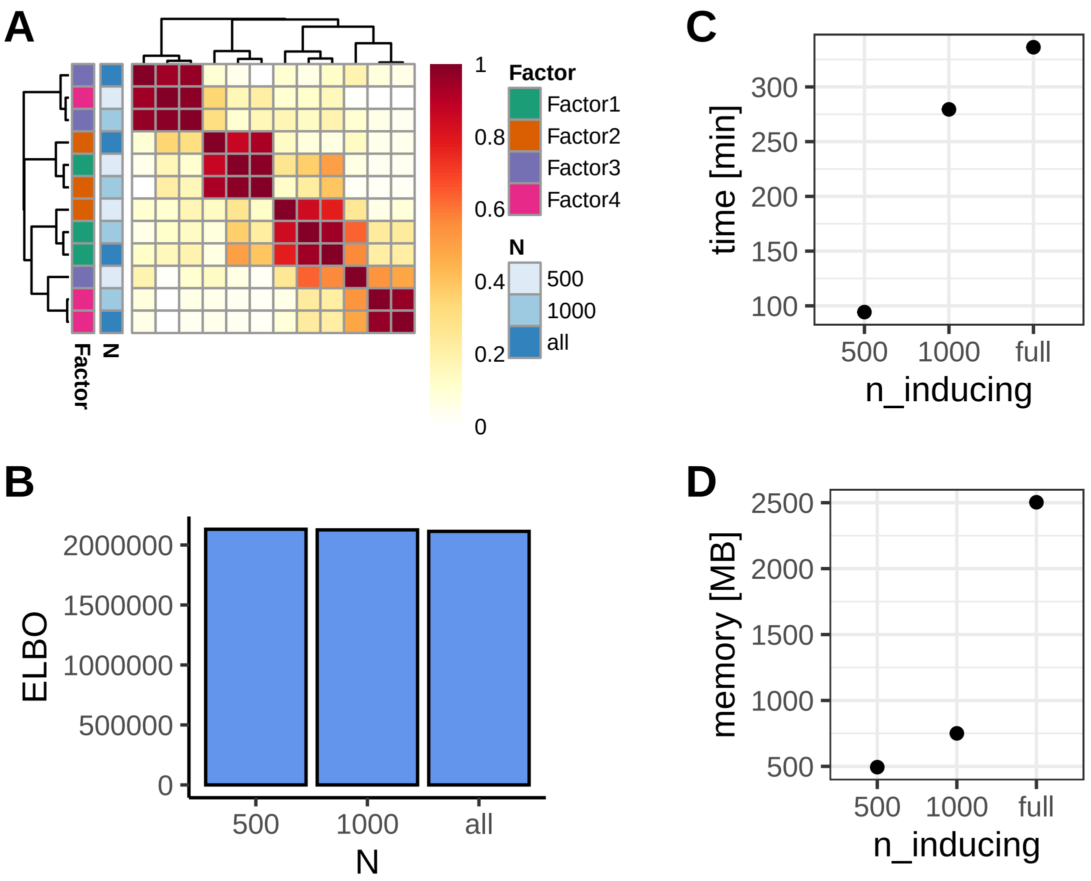


**Supp. Fig. 19: Evaluation of a sparse version of MEFISTO on spatial transcriptomics data**
MEFISTO was trained on the spatial transcriptomics data using varying number of inducing points (500, 1000) or on the full data (2696 spots). Shown is (A) the factor correlation between the resulting models, (B) the values of the evidence lower bound (ELBO) as well as (C) time and (D) memory requirements.
