## Supplementary Methods for "Identifying temporal and spatial patterns of variation from multi-modal data using MEFISTO"

**7 Comparison to related approaches** **17**

### 1 Introduction

MEFISTO provides an unsupervised approach to integrate multi-modal data with continuous structures among the samples, e.g. given by spatial or temporal relationships. Based on probabilistic factor analysis, it extends existing models to incorporate such dependencies and enable spatio-temporally informed dimensionality reduction as well as open up novel downstream analyses. MEFISTO builds upon on a recent framework for integration of multi-modal data (Multi-Omics Factor Analysis (MOFA)) [1, 2] that provides a sparse structured decomposition of the multi-modal input data to disentangle different sources of variation across data modalities and find a joint low-dimensional representation of the data in a common factor space. MEFISTO takes a functional view on this model in order to incorporate continuous covariates, while maintaining the ability to account for datasets consisting of diverse feature sets and sample groups. An important example for the use of MEFISTO are temporal or spatial data sets, where the covariate is given by time points or spatial coordinates. Here, it can disentangle patterns of variation that change smoothly along time or space from other sources of variation that are independent of time or space. In addition, it enables to interpolate and extrapolate to unseen or missing time points or locations as well as to cluster and align these patterns across multiple data sets. For an illustration of the method refer to **Figure 1** in the manuscript. In the following, we provide details on the model formulation and inference.

#### Mathematical notation

- Matrices are denoted with bold capital letters: $\mathbf{W}$
- Vectors are denoted with bold non-capital letters: $\mathbf{w}$. If the vector originates from a matrix, a single index will indicate the row that it comes from. If two indices are used, the first one corresponds to the row, the second one to the column and ':' denotes the entire row or column, e.g. $\mathbf{w}_i$ refers to the $i$th row and $\mathbf{w}_{:,j}$ refers to the $j$th column of the matrix $\mathbf{W}$.
- Scalars are denoted with non-bold and non-capital letters: $w$. If the scalar originates from a 1-dimensional array (a vector), a single subscript will indicate its position in the vector. If the scalar comes from a 2-dimensional array (a matrix), two indices will indicate its position in the array with the first indicating the row and the second indicating the column, e.g. $w_{i,j}$ refers to the value from the $i$th row and the $j$th column of the matrix $\mathbf{W}$ and $w_i$ to the $i$th value of the vector $\mathbf{w}$. For higher dimensional arrays (tensors) more than two indices are used accordingly.
- $\mathbf{0}_k$ is used to denote a zero vector of length $k$.
- $\mathbf{I}_k$ is used to denote the identity matrix of rank $k$.
- $\mathbb{E}_q[x]$ denotes the expectation of $x$ under the distribution $q$. If the distribution is clear from the context we will also use $\langle x \rangle$ to avoid cluttered notation.
- $\mathcal{N}(x \mid \mu, \sigma)$: $x$ follows a normal distribution with mean $\mu$ and variance $\sigma$.
- $\mathcal{N}(\mathbf{x} \mid \boldsymbol{\mu}, \boldsymbol{\Sigma})$: $\mathbf{x}$ follows a multivariate normal distribution with mean $\boldsymbol{\mu}$ and covariance matrix $\boldsymbol{\Sigma}$.
- $\mathcal{G}(x \mid a, b)$: $x$ follows a gamma distribution with shape and rate parameters $a$ and $b$.
- $\text{Beta}(x \mid a, b)$: $x$ follows a beta distribution with shape and rate parameters $a$ and $b$.
- $\text{Ber}(x|\theta)$: $x$ follows a Bernoulli distribution with parameter $\theta$.
- $\text{GP}(x|\mu, \kappa)$: $x$ follows a Gaussian process distribution with mean function $\mu$ and covariance function $\kappa$.
- $Tr(\mathbf{X})$: Trace of the matrix $\mathbf{X}$
- $\delta_{ij}$: Kronecker delta function taking the value of 1 if and only if $i = j$ and zero otherwise

#### 2 The MEFISTO model

##### 2.1 The underlying factor analysis model

The basis of MEFISTO is a probabilistic factor analysis model that provides a representation of a (high-dimensional) dataset in terms of a low-dimensional denoised representation (given by the factors). This representation can be used for visualisation and down-stream analysis (similar to principal components). Weight matrices provide a mapping from the low-dimensional factor space to the original feature space of the observed data and thereby can help to interpret patterns of variation captured by the factors. Recently, we proposed Multi-Omics Factor Analysis (MOFA, [1, 2]) as a multi-view generalisation of factors analysis building on the group factor analysis framework [3, 4]. MOFA finds a joint representation of $M$ data matrices (or views) $\mathbf{Y}^m \in \mathbb{R}^{N \times D_m}$ containing measurements on $N$ (common) samples and $D_m$ (view-specific) features in terms of $K$ joint factors and their view-specific weights. These views can for example be different omic modalities or distinct feature sets from a single omic modality (e.g. defined by genomic contexts). The basic underlying decomposition is given by

$$\mathbf{Y}^m = \mathbf{Z}\mathbf{W}^{mT} + \boldsymbol{\epsilon}^m, \tag{1}$$

where $\mathbf{Z} \in \mathbb{R}^{N \times K}$ contains the factor values, $\mathbf{W}^m \in \mathbb{R}^{D_m \times K}$ contains the weights that relate the factor values to the original data in view $m$ and $\boldsymbol{\epsilon}^m \in \mathbb{R}^{D_m}$ contains the residual variation that is not explained by the factors. Importantly, the structure of the data sets are encoded by prior distributions on the weights in the decomposition, which can encourage view- and feature-wise sparsity, thereby making it easier to pinpoint the views and features that underly the variation captured by a factor. This is achieved by a feature-wise spike-and-slab and view-wise automatic relevance determination (ARD) prior, which can be conveniently written as a combination of a Bernoulli random variable with view-wise and factor-wise probability ($\theta_k^m$) controlling the feature-wise sparsity per factor and a Gaussian random variable with view-wise and factor-wise precision ($\alpha_k^m$) controlling the view-wise sparsity per factor. The sparsity-controlling parameters are both learned and modelled by a Beta ($\theta_k^m$) or Gamma ($\alpha_k^m$) distribution. Please refer to [1] for details on the weight model.

The factors are modelled by a simple univariate prior, i.e.

$$p(z_{nk}) = \mathcal{N}\left(z_{nk} \,|\, 0, 1\right). \tag{2}$$

For details on the model we refer to the Supplementary Material of [1].

**Extension to a multi-group setting** More recently, the above model has been extended to data sets consisting of G distinct groups of samples $\mathbf{Y}^{m,g} \in \mathbb{R}^{N_g \times D_m}$ [2] . Here, the goal is to not only disentangle variation across views but to also across groups. For this, the method centers the features in each group to remove group-specific offsets and then learns group-specific factors, i.e.

$$\mathbf{Y}^{m,g} = \mathbf{Z}^g\mathbf{W}^{mT} + \boldsymbol{\epsilon}^{m,g}, \tag{3}$$

where $\mathbf{Z}^g \in \mathbb{R}^{N_g \times K}$ contains the factor values for group $g$. Again, the view structure of the data is encoded by prior distributions on the weights. In this multi-group setting the model furthermore employs similar prior distributions on the factors, which can additionally encourage group- and sample-wise sparsity. Here, the model still employs univariate priors, however now with group- and factor-wise parameters, that determine the activity of a factor per group:

$$p(z_{nk}^g) = \mathcal{N}\left(z_{nk} \,|\, 0, 1/\alpha_k^g\right). \tag{4}$$

Details are described in the Supplementary Material of [2].

**Limitations of these models** The above model formulations can incorporate discrete structures in the data, such as views and groups. However, they cannot easily be extended to data sets with continuous structure, which for example naturally occur in temporal or spatial data sets. In such data, continuous sample relationships are known and could guide the decomposition and inference of the factors. In particular, previous formulations of this model [1, 2] have used univariate priors for the factors, which provide a simple and flexible prior for inference but do not allow to model continuous sample relationships that should be reflected in the latent space. For this, we will in the following take a functional view on the factors and extend the above models to incorporate continuous covariates while maintaining the ability to incorporate the view- and group structure as well as feature-wise sparsity. We start with a description of the resulting model for a single group (Section 2.2), before extending it further to multiple sample groups in Section 2.3.

#### 2.2 A functional version of multi-omics factor analysis accounting for continuous covariates

In the following, we will assume that in addition to our data $\mathbf{Y}^m \in \mathbb{R}^{N\times D_m}$ we observe for each sample a continuous covariate $\mathbf{c}_n \in \mathbb{R}^C$. For instance, $\mathbf{c}_n$ could be a 1-dimensional covariate providing a time point for sample $n$ (e.g. age, developmental time, disease progression stage) or a 2-dimensional covariate providing the position of a sample in space (e.g. in a tissue, geographical or in a latent space).

To encode this additional information, MEFISTO uses a Gaussian process (GP) prior on the latent factors $\mathbf{z}_{:,k}$, $k = 1, \dots, K$, which are taken as realization of latent processes $f_k$ along the covariate

$$f_k \sim \mathrm{GP}(0, \kappa_k), \tag{5}$$

$$z_{nk} = f_k(\mathbf{c}_n) + \eta_{n,k} \tag{6}$$

$$\eta_{n,k} \sim \mathcal{N}(0, \zeta_k) \tag{7}$$

The properties of a latent process $k$, such as the smoothness along the covariate, is modelled by the covariance function $\kappa_k : \mathbb{R}^C \times \mathbb{R}^C \to \mathbb{R}$ of the Gaussian process, which is defined in terms of the covariate values. Broadly speaking, factor values for samples with similar covariates will have a high covariance, thereby encouraging smooth factors. By default, MEFISTO uses a squared exponential kernel with Euclidean distances to define a covariance function, which is given by

$$\kappa_k(\mathbf{c}_n, \mathbf{c}_{n'}) = s_k \exp\left(-\frac{||\mathbf{c}_n - \mathbf{c}_{n'}||_2^2}{2\ell_k^2}\right) \qquad \text{with } s_k = 1 - \zeta_k \tag{8}$$

with factor-wise parameters $\zeta_k, \ell_k$ that are learnt by optimizing them jointly with the other model components. The lengthscale parameter $\ell_k$ in the covariance function controls the speed at which the sample correlation decays along the covariate. The scale parameter $s_k = 1 - \zeta_k$ controls the proportion of smooth variation captured by the factor and is coupled to the variance of the noise $\eta$. This enables to distinguish factors of varying degree of smoothness as well as non-smooth factors, for which $\zeta_k = 1$ and where the covariance is diagonal, recovering the univariate prior of the original MOFA framework [1]. Extensions to other types of kernels and distance measures are straightforward and can be useful depending on the covariate and application, e.g. to detect periodic patterns. A general introduction to Gaussian processes can be found in [5].

#### 2.3 Modelling latent processes for multiple sample groups

As for the original MOFA model, we can further extend the functional model to datasets that consist of multiple groups of samples. For instance we might be given time course data from multiple individuals or species, where the individuals/species represent the groups and the time points the samples within each group. For this, we can use the same decomposition as in Section 2.1 for multiple groups but again with a functional view as well as additionally a continuous model of the relationships between group. In particular, we explicitly model the group-group correlation structure in the latent space by including the group information in the kernel function of the Gaussian process.

Denoting the covariate of sample $n$ in group $g$ by $\mathbf{c}_n^g \in \mathbb{R}^C$ and its factor value by $\mathbf{z}_n^g$ we define the $k$-th latent process as

$$f_k \sim \mathrm{GP}(0, \kappa_k), \tag{9}$$

$$\kappa_k(\mathbf{c}_n^g, \mathbf{c}_{n'}^{g'}) = s_k \exp\left(-\frac{||\mathbf{c}_n^g - \mathbf{c}_{n'}^{g'}||_2^2}{2\ell_k^2}\right) \kappa_k^G(g, g') \qquad \text{with } s_k = 1 - \zeta_k \tag{10}$$

$$z_{nk}^g = f_k(\mathbf{c}_n^g) + \eta_{n,k}^g \tag{11}$$

$$\eta_{n,k}^g \sim \mathcal{N}(0, \zeta_k) \tag{12}$$

The covariance function here combines the covariate kernel with a group-covariance matrix $(K_k^G)_{g,g'} = \kappa_k^G(g, g')$ which captures the relationships of the different groups on the latent process $k$. If one sets $(K_k^G)_{gg'} = 1 \ \forall g, g'$, this corresponds to a concatenation of the samples from all groups and thereby encourages the model to learn factor values that show very similar profiles along the covariates in each sample group by having a-priori a perfect correlation between factor values from different groups at the same value of the covariate. To account for group differences and learn $K_k^G$ in a data-driven manner we use a low-rank approximation of $K_k^G$ following prior work in the area of multi-task Gaussian process regression [6], i.e. we define

$$\tilde{K}_k^G = \sum_{r=1}^{R} \mathbf{x}_r^{(k)} \mathbf{x}_r^{(k)T} + \sigma_k^2 \mathbf{I}_G \quad \text{with } x_r^{(k)} \in \mathbb{R}^G \tag{13}$$

and set $K_G$ to be the correlation matrix corresponding to this covariance matrix $\tilde{K}_k^G$, which ensures all values lie between -1 and 1. The value of $R$ controls the rank of the approximation and is set to 1 or 2 depending on the number of groups. An illustration of the resulting covariance structures for two groups is given in Figure 1.

Note that a joint modelling of distinct sample groups is only reasonable if we have an accurate correspondence between the covariates across groups. However, in some setting this might not be the case (e.g. correspondences of developmental stages between species or disease progression time courses between patients). To address this problem and enable simultaneously aligning and factorizing the data we introduce an alignment procedure in Section 5.

In the following, we will drop the group index from the notation, where it is not required, and use $\mathbf{z}_n$ to denote the concatenation of the factor values across groups. We use $N_g$ to denote the number of samples in group $g$ and $N$ to denote the total number of sample, i.e. $N = \sum_{g=1}^{G} N_g$.

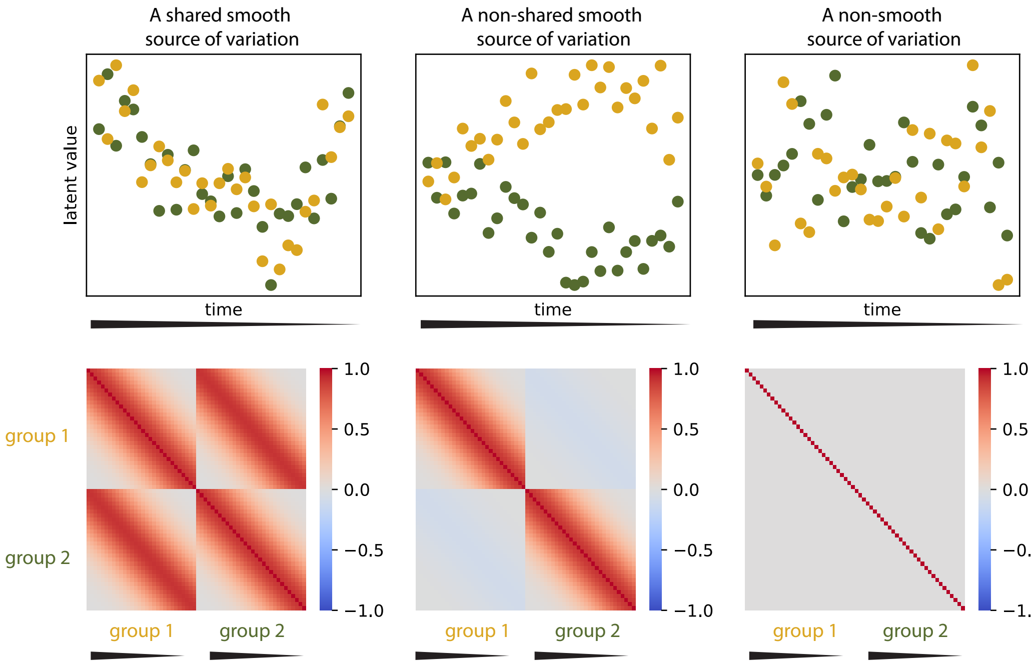

Figure 1: Example illustrating the group and continuous-covariance structure in a setting with a smooth latent process shared between groups (left), a smooth latent process that is distinct between groups (middle) and a non-smooth latent process (right).

#### 2.4 A full model specification

Taken together, the above results in the following model specification for MEFISTO :

$$
\begin{aligned}
p(y^m_{nd}|\mathbf{Z}, \mathbf{W}, \boldsymbol{\tau}) &= \mathcal{N}\left(y^m_{nd}|\sum_k w^m_{kd} z^g_{nk}, 1/\tau^{m,g}_d\right) && (14)\\
p(f_k) &= \mathrm{GP}(f_k \,|\, 0, \kappa_k) && (15)\\
&\quad \text{with} \quad \kappa_k(\mathbf{c}^g_n, \mathbf{c}^{g'}_{n'}) = (1-\zeta_k)\exp\left(-\frac{||\mathbf{c}^g_n - \mathbf{c}^{g'}_{n'}||^2_2}{2\ell^2_k}\right)\kappa^G_k(g, g') && (16)\\
p(z^g_{nk}|f_k) &= f_k(\mathbf{c}^g_n) + \eta^g_{n,k} && (17)\\
p(\eta^g_{n,k}) &= \mathcal{N}(0, \zeta_k) && (18)\\
w^m_{kd} &= \hat{w}^m_{kd} s^m_{kd} && (19)\\
&\quad \text{with} \quad p(\hat{w}^m_{kd}, s^m_{kd}|\theta^m_k, \alpha^m_k) = \mathcal{N}(\hat{w}^m_{kd} \,|\, 0, 1/\alpha^m_k)\,\mathrm{Ber}(s^m_{kd} \,|\, \theta^m_k) && (20)\\
p(\theta^m_k) &= \mathrm{Beta}\left(\theta^m_k \,|\, a^\theta_0, b^\theta_0\right) && (21)\\
p(\alpha^m_k) &= \mathcal{G}\left(\alpha^m_k \,|\, a^\alpha_0, b^\alpha_0\right) && (22)\\
p(\tau^{m,g}_d) &= \Gamma(a^\tau_0, b^\tau_0). && (23)
\end{aligned}
$$

Here, the factors are modelled as described above and the view-wise weights are modelled as in [1], recapitulated in Section 2.1, with fixed hyper-parameters $a^\theta_0, b^\theta_0 = 1$ and $a^\tau_0, b^\tau_0, a^\alpha_0, b^\alpha_0 = 0.001$ to obtain uninformative priors. The hyperparameters of the covariance function $\kappa_k$, i.e. $\zeta_k$, $\ell_k$, $\mathbf{x}^{(k)}$ and $\sigma_k$ are learnt during training and we describe details in Section 3.2. In the following we will often use the marginal distribution for the values $\mathbf{z}_k$ on the $k$-th factor, which is given by

$$
p(\mathbf{z}_{:,k}) = \mathcal{N}\left(\mathbf{z}_{:,k}|0, \boldsymbol{\Sigma}_k\right) \quad \text{with} \quad (\boldsymbol{\Sigma}_k)_{nn'} = (1-\zeta_k)\exp\left(-\frac{||\mathbf{c}^g_n - \mathbf{c}^{g'}_{n'}||^2}{2\ell^2_k}\right)\kappa^G_k(g, g') + \zeta_k \delta_{n,n'}, \qquad (24)
$$

where the factor values $\mathbf{z}_{:,k}$ are concatenated along the sample axis for all groups and $g, g'$ denote the groups corresponding to samples $n, n'$.

While we formulated the model with Gaussian noise, other noise models can be used as implemented in the MOFA framework (see [1, 2] for details). The complete model is illustrated in Figure 2.

### 3 Inference in the MEFISTO model

While large parts of the original variational inference framework from MOFA [1, 2] can be re-used in order to approximate the posterior distribution from the model, we need to develop a new approach for the inference of the factors $\mathbf{Z}$. In addition, simultaneous to the updates of the main model components the hyperparameters of the Gaussian processes for each factor need to be found. Due to the introduction of a Gaussian process prior on the factors the evidence lower bound is no longer decomposable in the samples and stochastic inference as implemented in [2] does not easily generalize. Here, we will first adapt the full inference framework to a model with Gaussian process prior and a multivariate variational distribution. In section 4, we will provide details on a sparse approximation of the Gaussian process using inducing points that can be used to achieve a better scalability in terms of computation time and memory in the presence of many samples.

#### 3.1 Short introduction to variational Bayes

Given a probabilistic model with observed variables $\mathbf{Y}$ and latent variables $\mathbf{X}$, we are interested in finding the posterior distribution of the latent variables given the observations $p(\mathbf{X}|\mathbf{Y}) = \frac{p(\mathbf{Y},\mathbf{X})}{\int_x p(\mathbf{Y},\mathbf{X})}$. However, often the posterior cannot be derived in an analytical form and therefore approximations are necessary. For this purpose, different approaches exist: sampling-based methods (Markov Chain Monte Carlo) and approaches that recast the problem of finding the posterior as an optimization problem. Variational inference takes the latter approach and can be much faster than sampling based inference. The key idea is to approximate the true posterior $p(\mathbf{X}|\mathbf{Y})$ by a more tractable variational distribution $q(\mathbf{X})$

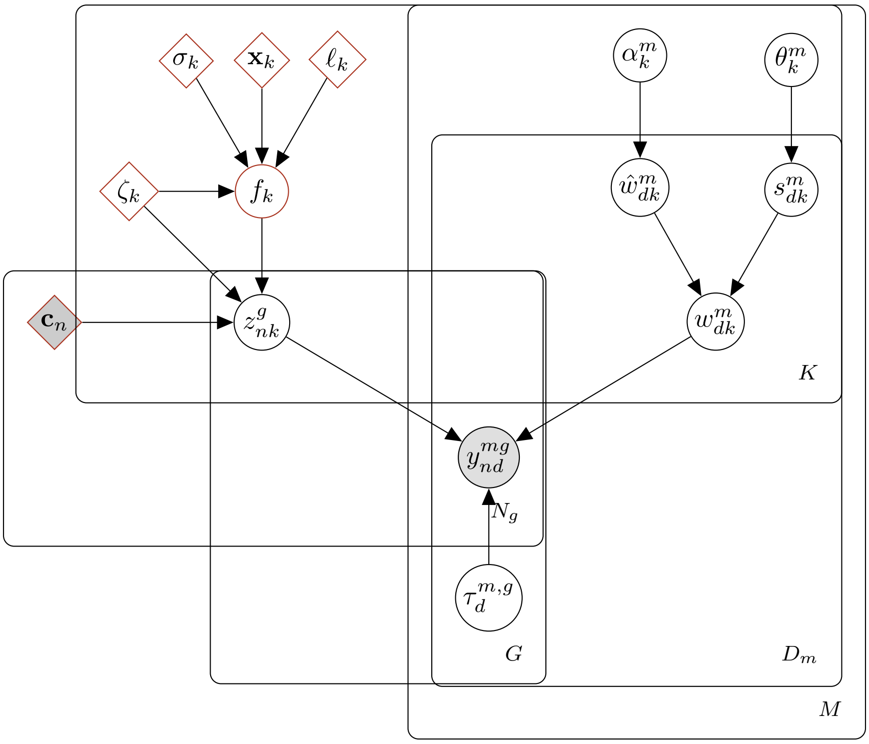

Figure 2: Illustration of the MEFISTO model. Observed nodes are colored in grey, unobserved nodes in white. Hyperparameters and non-probabilistic nodes are shown as a rectangle. For comparison to MOFA, nodes that are added in MEFISTO are marked by brown borders.

and minimizing the 'distance' (measured by the Kullback-Leibler (KL) divergence) of this variational distribution to the true posterior:

$$\mathrm{KL}(q(\mathbf{X})||p(\mathbf{X}|\mathbf{Y})) = -\int_x q(\mathbf{X}) \log \frac{p(\mathbf{X}|\mathbf{Y})}{q(\mathbf{X})} \tag{25}$$

As this term still contains the intractable posterior, this does not yet simplify the problem. However, one can show that instead of minimizing the KL divergence it is possible to maximize a quantity $\mathcal{L}(q(\mathbf{X}))$ called Evidence Lower Bound (ELBO) thanks to the following decomposition, where the left-hand side does not depend on the variational distribution:

$$\log p(\mathbf{Y}) = \mathrm{KL}(q(\mathbf{X})||p(\mathbf{X}|\mathbf{Y})) + \mathcal{L}(q(\mathbf{X})) \tag{26}$$

Here, the ELBO is given as

$$\begin{aligned}\mathcal{L}(q(\mathbf{X})) &= \int_x q(\mathbf{X}) \Big( \log \frac{p(\mathbf{X}|\mathbf{Y})}{q(\mathbf{X})} + \log p(\mathbf{Y}) \Big) d\mathbf{X} \\ &= \mathbb{E}_q[\log p(\mathbf{X}, \mathbf{Y})] - \mathbb{E}_q[\log q(\mathbf{X})] \\ &= \mathbb{E}_q[\log p(\mathbf{Y}|\mathbf{X})] + (\mathbb{E}_q[\log p(\mathbf{X})] - \mathbb{E}_q[\log q(\mathbf{X})]).\end{aligned} \tag{27}$$

Note that this term only depends on the known joint model density and the variational distribution, which we restrict to members of a suitable family of distributions, and hence can be directly optimized, recasting the problem of finding the posterior as an optimization problem. Crucial is the choice of the class of variational distribution to consider. Common approaches include parametric assumptions (paired with gradient-based optimization of the ELBO) or free-form approaches often taking a mean-field assumption, where we assume $q(\mathbf{X}) = \prod_{i=1}^M q_i(\mathbf{x}_i)$, resulting in explicit update equations in conjugate models given by

$$\log q_i^*(\mathbf{x}_i) = \mathbb{E}_{-i}[\log p(\mathbf{Y}, \mathbf{X})] + \text{const.} \tag{28}$$

A more detailed introduction to variational methods can be found in [7–9].

#### 3.2 Updates for the MEFISTO model

Due to the dependencies between the samples that are given by the continuous covariates, MEFISTO does not follow the complete mean-field assumption of MOFA and MOFA+ [1, 2], where a full factorization into univariate variational distributions was assumed. Instead, we make the following assumption on the factorization of the variational distribution and from this derive the updates as given by (28).

$$q(\mathbf{X}) = \prod_{k=1}^{K} q(\mathbf{z}_{:,k}) \prod_{m=1}^{M} \prod_{d=1}^{D_m} \prod_{k=1}^{K} q(\hat{w}_{kd}^m, s_{kd}^m) \prod_{m=1}^{M} \prod_{k=1}^{K} q(\alpha_k^m) \prod_{m=1}^{M} \prod_{k=1}^{K} q(\theta_k^m) \prod_{m=1}^{M} \prod_{d=1}^{D_m} q(\tau_d^m). \tag{29}$$

By this factorization, we neglect posterior correlations between distinct factors in the variational distribution but importantly make use of a multivariate distribution to model correlation between values on the same factor.

In addition, the covariance structure of the prior $p(\mathbf{z}_{:,k})$ is determined by the GP hyperparameters, that are optimized alongside with the variational updates by maximising the ELBO. We iteratively optimize first every component $i$ of the variational distribution

$$q_i^*(\mathbf{x}_i) = \arg\max_{q_i} \mathcal{L}_{\ell,\mathbf{x},\zeta,\sigma}(q), \tag{30}$$

followed by an optimization of the hyperparameters

$$\ell, \mathbf{x}, \zeta, \sigma = \arg\max_{\ell,\mathbf{x},\zeta,\sigma} \mathcal{L}_{\ell,\mathbf{x},\zeta,\sigma}(q). \tag{31}$$

In the following, we provide updates for all components in the variational distribution and the Gaussian process hyperparameters.

##### 3.2.1 Updates of the latent factors

For every factor $k$ we have the following prior distribution $p(\mathbf{z}_{:,k})$:

$$p(\mathbf{z}_{:,k}) = \mathcal{N}(\mathbf{z}_{:,k}|0, \mathbf{\Sigma}_k) \tag{32}$$

The latent factors $\mathbf{z}_{:,k}$ are modelled with a multivariate variational distribution that leads to the following updates: For each factor $k$ we have

$$q(\mathbf{z}_{:,k}) = \mathcal{N}(\mathbf{z}_{:,k}, |\boldsymbol{\mu}_k, \mathbf{A}_k), \tag{33}$$

where

$$\begin{aligned} \mathbf{A}_k &= \left( \text{diag}(\sum_{d,m} \langle w_{dk}^{(m)2} \rangle \langle \tau_d^{(m)} \rangle) + \mathbf{\Sigma}_k^{-1} \right)^{-1} \\ \mathbf{A}_k^{-1}\boldsymbol{\mu} &= \left( \sum_{m,d} \langle \tau_d^{(m)} \rangle \langle w_{dk}^{(m)} \rangle \left( y_{nd}^{(m)} - \sum_{l \neq k} \langle w_{dl}^{(m)} \rangle \langle z_{nl} \rangle \right) \right)_{n=1,\ldots,N} \end{aligned} \tag{34}$$

##### 3.2.2 Updates of the Gaussian process hyperparameters

For the Gaussian processes we need to optimize the hyperparameters given by the lengthscale $\ell_k$ and scale $s_k = 1 - \zeta_k$ per factor. With multiple groups, additionally the hyperparameters $\mathbf{x}_k$ and $\sigma_k$ need to be found. Due to the dependence of the prior and variational distribution of $\mathbf{Z}$ on $\mathbf{\Sigma}_k$ the value of the ELBO depends on the hyperparametes via

$$\mathcal{L}_{\ell,\mathbf{x},\zeta,\sigma}(q) = \mathbb{E}_q[\log p(\mathbf{Y}|\mathbf{X})] + (\mathbb{E}_q[\log p_{\ell,\mathbf{x},\zeta,\sigma}(\mathbf{X})] - \mathbb{E}_q[\log q(\mathbf{X})]), \tag{35}$$

where in the second part only the terms involving $\mathbf{Z}$ depend on the hyperparameters, i.e. $\mathbb{E}_q[\log p_{\ell,\mathbf{x},\zeta,\sigma}(\mathbf{Z})] - \mathbb{E}_q[\log q((\mathbf{Z})]$. These are given by

$$\mathbb{E}_q[\log p_{\ell,\mathbf{x},\zeta,\sigma}(\mathbf{Z})] = \sum_{k=1}^{K} \mathbb{E}_q[\log p_{\ell,\mathbf{x},\zeta,\sigma}(\mathbf{z}_{:,k})] \tag{36}$$

$$= \sum_{k=1}^{K} \left( \frac{1}{2} \log\det(\mathbf{\Sigma}_k^{-1}) - \frac{1}{2} \langle \mathbf{z}_{:,k}^T \mathbf{\Sigma}_k^{-1} \mathbf{z}_{:,k} \rangle - \frac{N}{2} \log(2\pi) \right) \tag{37}$$

$$\mathbb{E}_q[\log q((\mathbf{Z})] = -\sum_{k=1}^{K} \frac{1}{2} \log\det(\mathbf{A}_k) - \frac{KN}{2} - \frac{KN}{2} \log(2\pi). \tag{38}$$

As this decomposes in $k$, the optimization of the lengthscale and scale hyperparametere $\ell = (\ell_1, \dots, \ell_K)$, $\zeta = (\zeta_1, \dots, \zeta_K)$, $\mathbf{x} = (\mathbf{x}_1, \dots, \mathbf{x}_K)$ and $\sigma = (\sigma_1, \dots, \sigma_K)$ can be performed independent for each factor given the variational distributions of the current iteration.

$$\ell_k, \zeta_k, \mathbf{x}_k, \sigma_k = \arg \max_{\ell_k, \zeta_k, \mathbf{x}_k, \sigma_k} \frac{1}{2} \log\det(\mathbf{\Sigma}_k^{-1}) - \frac{1}{2} \langle \mathbf{z}_{:,k}^T \mathbf{\Sigma}_k^{-1} \mathbf{z}_{:,k} \rangle + \text{const.} \tag{39}$$

$$= \arg \max_{\ell_k, \zeta_k, \mathbf{x}_k, \sigma_k} \frac{1}{2} \log\det(\mathbf{\Sigma}_k^{-1}) - \frac{1}{2} \mathrm{tr}(\mathbf{\Sigma}_k^{-1} \mathbf{A}_k) - \frac{1}{2} \boldsymbol{\mu}_k^T \mathbf{\Sigma}_k^{-1} \boldsymbol{\mu}_k + \text{const.}, \tag{40}$$

where $\mathbf{\Sigma}_k$ is defined via the hyperparameters as given in Equation (24). This maximization is performed using a grid search for possible values of $\ell_k$ ranging from half the minimal distance to twice the maximal distance between samples (in terms of the covariate $\mathbf{C}$) following [10, 11] and zero. For each grid point the optimal $\zeta_k, \sigma_k$ and $\mathbf{x}_k$ are found by optimization of the objective function in equation (39) using L-BFGS-B in $(0, 1)$ (for $\zeta_k$ and $\sigma_k$) and $[-1, 1]$ (for $\mathbf{x}_k$). As the model is not identifiable for some configurations of the hyperparameters we set $\zeta_k = 1, \sigma_k = 1, \mathbf{x}_k = \mathbf{0}$ and $\ell_k = 0$ if either the lengthscale is zero or the scale $1 - \zeta_k$ is zero, which corresponds to an unstructured prior with no group or covariate kernel.

If only a single sample group is present or the samples have Kronecker structure, i.e. for each value of the covariate one sample was observed in each group, we can use a spectral decomposition of the kernel matrices in order to avoid re-calculating the inverse of the covariance matrix and its determinant and enable a more efficient inference than a naive optimization. In this case, we denote with $T = N_g$ the (common) number of samples per group with covariate values $\mathbf{c}_t,\ t = 1, \dots, T$.

$$\mathbf{\Sigma}_k = (1 - \zeta_k) \mathbf{K}_{\mathbf{x}_k, \sigma_k}^{(G)} \otimes \mathbf{K}_{\ell_k}^{(C)} + \zeta_k \mathbf{I}_{TG} \tag{41}$$

$$\text{with} \quad \mathbf{K}_{\ell_k}^{(C)} = \left( \exp\left( -\frac{||\mathbf{c}_t - \mathbf{c}_{t'}||^2}{2\ell_k^2} \right) \right)_{t,t'=1,\dots,T} \tag{42}$$

$$\tilde{\mathbf{K}}_k^{(G)} = \sum_{r=1}^{R} \mathbf{x}_r^{(k)} \mathbf{x}_r^{(k)T} + \sigma_k^2 \mathbf{I}_G \tag{43}$$

As $\mathbf{K}^{(G)}$ (defined as the correlation-matrix corresponding to $\tilde{\mathbf{K}}_k^{(G)}$) and $\mathbf{K}^{(C)}$ are symmetric, we can decompose them into a diagonal matrix $\mathbf{D}$ and a orthogonal matrix $\mathbf{V}$ with $\mathbf{V}^T \mathbf{V} = \mathbf{I}$

$$\mathbf{K}_{\ell_k}^{(C)} = \mathbf{V}_{\ell_k}^{(C)} \mathbf{D}_{\ell_k}^{(C)} \mathbf{V}_{\ell_k}^{(C)T}. \tag{44}$$

and

$$\mathbf{K}_{\mathbf{x}_k, \sigma_k}^{(G)} = \mathbf{V}_{\mathbf{x}_k, \sigma_k}^{(G)} \mathbf{D}_{\mathbf{x}_k, \sigma_k}^{(G)} \mathbf{V}_{\mathbf{x}_k, \sigma_k}^{(G)T}. \tag{45}$$

Using this we can calculate $\mathbf{\Sigma}_k$, its inverse and determinant as

$$\mathbf{\Sigma}_k = (1 - \zeta_k)(\mathbf{V}^{(G)} \otimes \mathbf{V}^{(C)})(\mathbf{D}^{(G)} \otimes \mathbf{D}^{(C)} + \frac{\zeta_k}{1 - \zeta_k} \mathbf{I}_G \otimes \mathbf{I}_T)(\mathbf{V}^{(G)T} \otimes \mathbf{V}^{(C)T}) \tag{46}$$

$$\mathbf{\Sigma}_k^{-1} = \frac{1}{1 - \zeta_k} (\mathbf{V}^{(G)} \otimes \mathbf{V}^{(C)})(\mathbf{D}^{(G)} \otimes \mathbf{D}^{(C)} + \frac{\zeta_k}{1 - \zeta_k} \mathbf{I}_G \otimes \mathbf{I}_T)^{-1} (\mathbf{V}^{(G)T} \otimes \mathbf{V}^{(C)T}) \tag{47}$$

$$\log\det \mathbf{\Sigma}_k^{-1} = -TG \log(1 - \zeta_k) - \log\det(\mathbf{D}^{(G)} \otimes \mathbf{D}^{(C)} + \frac{\zeta_k}{1 - \zeta_k} \mathbf{I}_G \otimes \mathbf{I}_T) \tag{48}$$

as $\mathbf{V}^{(C)}, \mathbf{V}^{(G)}$ are orthogonal. This can be evaluated efficiently as it only requires the inversion of a diagonal matrix.

###### 3.2.3 Updates for the weights and noise terms

The remaining model updates are analogous to the MOFA model with univariate prior on $\mathbf{Z}$ and are reproduced below from [2] for completeness.

**Sparse weights (with spike-and-slab prior)** For every view $m$, feature $d$ and factor $k$:

Prior distribution $p(\hat{w}_{kd}^m, s_{kd}^m)$:

$$p(\hat{w}_{kd}^m, s_{kd}^m) = \mathcal{N}(\hat{w}_{kd}^m \,|\, 0,\ 1/\alpha_k^m)\,\mathrm{Ber}(s_{kd}^m \,|\, \theta_k^m) \tag{49}$$

Variational distribution $q(\hat{w}_{kd}^m, s_{kd}^m)$:

Update for $q(s_{kd}^m)$:

$$q(s_{kd}^m) = \mathrm{Ber}(s_{kd}^m | \gamma_{kd}^m) \tag{50}$$

with

$$\begin{aligned}
\gamma_{kd}^m &= \frac{1}{1+\exp(-\lambda_{kd}^m)} \\
\lambda_{kd}^m &= \langle \log \frac{\theta}{1-\theta} \rangle + 0.5 \log \frac{\langle \alpha_k^m \rangle}{\langle \tau_d^m \rangle} - 0.5 \log \left( \sum_{n=1}^{N} \langle (z_{nk})^2 \rangle + \frac{\langle \alpha_k^m \rangle}{\langle \tau_d^m \rangle} \right) \\
&+ \frac{\langle \tau_d^m \rangle}{2} \frac{\left( \sum_{n=1}^N y_{nd}^m \langle z_{nk} \rangle - \sum_{j \neq k} \langle s_{jd}^m \hat{w}_{jd}^m \rangle \sum_{n=1}^N \langle z_{nk} \rangle \langle z_{nj} \rangle \right)^2}{\sum_{n=1}^N \langle (z_{nk})^2 \rangle + \frac{\langle \alpha_k^m \rangle}{\langle \tau_d^m \rangle}}
\end{aligned} \tag{51}$$

Update for $q(\hat{w}_{kd}^m | s_{kd}^m)$:

$$\begin{aligned}
q(\hat{w}_{kd}^m | s_{kd}^m = 0) &= \mathcal{N}\left(\hat{w}_{kd}^m | 0, 1/\alpha_k^m\right) \\
q(\hat{w}_{kd}^m | s_{kd}^m = 1) &= \mathcal{N}\left(\hat{w}_{kd}^m \middle| \mu_{w_{kd}^m}, \boldsymbol{\Sigma}_{w_{kd}^m}^2\right)
\end{aligned} \tag{52}$$

with

$$\begin{aligned}
\mu_{w_{kd}^m} &= \frac{\sum_{n=1}^N y_{nd}^m \langle z_{nk} \rangle - \sum_{j \neq k} \langle s_{jd}^m \hat{w}_{jd}^m \rangle \sum_{n=1}^N \langle z_{nk} \rangle \langle z_{nj} \rangle}{\sum_{n=1}^N \langle (z_{nk})^2 \rangle + \frac{\langle \alpha_k^m \rangle}{\langle \tau_d^m \rangle}} \\
\boldsymbol{\Sigma}_{w_{kd}^m}^2 &= \frac{\langle \tau_d^m \rangle^{-1}}{\sum_{n=1}^N \langle (z_{nk})^2 \rangle + \frac{\langle \alpha_k^m \rangle}{\langle \tau_d^m \rangle}}
\end{aligned} \tag{53}$$

**ARD precision of the weights** For every view $m$ and factor $k$:

Prior distribution $p(\alpha_k^m)$:

$$p(\alpha_k^m) = \mathcal{G}\left(\alpha_k^m \,|\, a_0^\alpha, b_0^\alpha\right)$$

Variational distribution $q(\alpha_k^m)$:

$$q(\alpha_k^m) = \mathcal{G}\left(\alpha_k^m \,|\, \hat{a}_{mk}^\alpha, \hat{b}_{mk}^\alpha\right) \tag{54}$$

with

$$\begin{aligned}
\hat{a}_{mk}^\alpha &= a_0^\alpha + \frac{D_m}{2} \\
\hat{b}_{mk}^\alpha &= b_0^\alpha + \frac{\sum_{d=1}^{D_m} \langle (\hat{w}_{kd}^m)^2 \rangle}{2}
\end{aligned} \tag{55}$$

**Sparsity parameter of the weights** For every view $m$ and factor $k$:

Prior distribution:

$$p(\theta_k^m) = \text{Beta}\left(\theta_k^m \mid a_0^\theta, b_0^\theta\right)$$

Variational distribution:

$$q(\theta_k^m) = \text{Beta}\left(\theta_k^m \mid \hat{a}_{mk}^\theta, \hat{b}_{mk}^\theta\right) \tag{56}$$

with

$$\begin{aligned} \hat{a}_{mk}^\theta &= \sum_{d=1}^{D_m} \langle s_{kd}^m \rangle + a_0^\theta \\ \hat{b}_{mk}^\theta &= b_0^\theta - \sum_{d=1}^{D_m} \langle s_{kd}^m \rangle + D_m \end{aligned} \tag{57}$$

**Noise (Gaussian)** For every view $m$ and feature $d$:

Prior distribution $p(\tau_d^m)$:

$$p(\tau_d^g) = \mathcal{G}\left(\tau_d^m \mid a_0^\tau, b_0^\tau\right),$$

Variational distribution $q(\tau_d^m)$:

$$q(\tau_d^m) = \mathcal{G}\left(\tau_d^m \mid \hat{a}_d^m, \hat{b}_d^m\right) \tag{58}$$

with

$$\begin{aligned} \hat{a}_d^m &= a_0^\tau + \frac{N}{2} \\ \hat{b}_d^m &= b_0^\tau + \frac{1}{2} \sum_{n=1}^{N} \langle \left( y_{nd}^m - \sum_k^K w_{kd}^m z_{nk} \right)^2 \rangle \end{aligned} \tag{59}$$

#### 3.3 Evidence lower bound

As described above, the evidence lower bound is composed of a log likelihood term and the Kullback Leibler-divergence between the prior and variational distribution of each unobserved model component ($\text{KL}\left(q(\mathbf{X}) || p(X)\right) = \mathbb{E}_q(q(\mathbf{X})) - \mathbb{E}_q(p(\mathbf{X}))$.).

$$\mathcal{L} = \mathbb{E}_q[\log p(\mathbf{Y}|\mathbf{X})] + \left(\mathbb{E}_q[\log p(\mathbf{X})] - \mathbb{E}_q[\log q(\mathbf{X})]\right) \tag{60}$$

##### 3.3.1 Contribution of the latent factor component

For a factor $k$ the term originating from the prior is given by:

$$\langle \log p(\mathbf{z}_{:,k}) \rangle = \frac{1}{2} \log \det(\mathbf{\Sigma}_k^{-1}) - \frac{1}{2} \langle \mathbf{z}_{:,k}^T \mathbf{\Sigma}_k^{-1} \mathbf{z}_{:,k} \rangle - \frac{N}{2} \log(2\pi) \tag{61}$$

The variational distribution $q(\mathbf{z}_{:,k})$ yields the following term for a factor $k$:

$$\langle \log q(\mathbf{z}_{:,k}) \rangle = -\frac{1}{2} \log \det(\mathbf{A}_k) - \frac{N}{2} - \frac{N}{2} \log(2\pi) \tag{62}$$

##### 3.3.2 Contributions of the likelihood term and the remaining components

The remaining terms of the ELBO are analogous to the MOFA model with univariate prior on Z and are reproduced below from [2] for completeness.

**Log likelihood term** Assuming a Gaussian likelihood:

$$\begin{aligned} \mathbb{E}_{q(X)} \log P(Y|X) = &- \sum_{m=1}^{M} \frac{ND_m}{2} \log(2\pi) + \frac{N}{2} \sum_{m=1}^{M} \sum_{d=1}^{D_m} \langle \log(\tau_d^m) \rangle \\ &- \sum_{m=1}^{M} \sum_{d=1}^{D_m} \frac{\langle \tau_d^m \rangle}{2} \sum_{n=1}^{N} \left( y_{nd}^m - \sum_{k=1}^{K} \langle s_{kd}^m \hat{w}_{kd}^m \rangle \langle z_{nk} \rangle \right)^2 \end{aligned} \tag{63}$$

**KL divergence terms**

* *Sparse weights*

$$\mathbb{E}_q[\log p(\hat{W}, S)] = -\sum_{m=1}^{M} \frac{KD_m}{2}\log(2\pi) + \sum_{m=1}^{M}\frac{D_m}{2}\sum_{k=1}^{K}\log(\alpha_k^m) - \sum_{m=1}^{M}\frac{\alpha_k^m}{2}\sum_{d=1}^{D_m}\sum_{k=1}^{K}\langle(\hat{w}_{kd}^m)^2\rangle$$
$$+ \langle\log(\theta)\rangle\sum_{m=1}^{M}\sum_{d=1}^{D_m}\sum_{k=1}^{K}\langle s_{kd}^m\rangle + \langle\log(1-\theta)\rangle\sum_{m=1}^{M}\sum_{d=1}^{D_m}\sum_{k=1}^{K}(1-\langle s_{kd}^m\rangle) \tag{64}$$

$$\mathbb{E}_q[\log q(\hat{W}, S)] = -\sum_{m=1}^{M}\frac{KD_m}{2}\log(2\pi) + \frac{1}{2}\sum_{m=1}^{M}\sum_{d=1}^{D_m}\sum_{k=1}^{K}\log(\langle s_{kd}^m\rangle\mathbf{\Sigma}_{w_{kd}^m}^2 + (1-\langle s_{kd}^m\rangle)/\alpha_k^m)$$
$$+\sum_{m=1}^{M}\sum_{d=1}^{D_m}\sum_{k=1}^{K}(1-\langle s_{kd}^m\rangle)\log(1-\langle s_{kd}^m\rangle) - \langle s_{kd}^m\rangle\log\langle s_{kd}^m\rangle \tag{65}$$

* *ARD precision for the weights*

$$\mathbb{E}_q[\log p(\boldsymbol{\alpha})] = \sum_{m=1}^{M}\sum_{k=1}^{K}\Big(a_0^\alpha\log b_0^\alpha + (a_0^\alpha - 1)\langle\log\alpha_k\rangle - b_0^\alpha\langle\alpha_k\rangle - \log\Gamma(a_0^\alpha)\Big)$$
$$\mathbb{E}_q[\log q(\boldsymbol{\alpha})] = \sum_{m=1}^{M}\sum_{k=1}^{K}\Big(\hat{a}_k^\alpha\log\hat{b}_k^\alpha + (\hat{a}_k^\alpha - 1)\langle\log\alpha_k\rangle - \hat{b}_k^\alpha\langle\alpha_k\rangle - \log\Gamma(\hat{a}_k^\alpha)\Big) \tag{66}$$

* *Sparsity parameter of the weights*

$$\mathbb{E}_q\left[\log p(\boldsymbol{\theta})\right] = \sum_{m=1}^{M}\sum_{k=1}^{K}\sum_{d=1}^{D_m}\big((a_0 - 1)\times\langle\log(\pi_{d,k}^m)\rangle + (b_0 - 1)\langle\log(1-\pi_{d,k}^m)\rangle - \log(\mathrm{B}(a_0, b_0))\big)$$
$$\mathbb{E}_q\left[\log q(\boldsymbol{\theta})\right] = \sum_{m=1}^{M}\sum_{k=1}^{K}\sum_{d=1}^{D_m}\big((a_{k,d}^m - 1)\times\langle\log(\pi_{d,k}^m)\rangle + (b_{k,d}^m - 1)\langle\log(1-\pi_{d,k}^m)\rangle - \log(\mathrm{B}(a_{k,d}^m, b_{k,d}^m))\big) \tag{67}$$

* *Noise*

$$\mathbb{E}_q[\log p(\boldsymbol{\tau})] = \sum_{m=1}^{M} D_m a_0^\tau\log b_0^\tau + \sum_{m=1}^{M}\sum_{d=1}^{Dm}(a_0^\tau - 1)\langle\log\tau_d^m\rangle - \sum_{m=1}^{M}\sum_{d=1}^{Dm} b_0^\tau\langle\tau_d^m\rangle - \sum_{m=1}^{M} D_m\log\Gamma(a_0^\tau)$$
$$\mathbb{E}_q[\log q(\boldsymbol{\tau})] = \sum_{m=1}^{M}\sum_{d=1}^{D_m}\Big(\hat{a}_{dm}^\tau\log\hat{b}_{dm}^\tau + (\hat{a}_{dm}^\tau - 1)\langle\log\tau_d^m\rangle - \hat{b}_{dm}^\tau\langle\tau_d^m\rangle - \log\Gamma(\hat{a}_{dm}^\tau)\Big) \tag{68}$$

#### 3.4 Notes on the complexity

Due to the appearance of the sample covariance matrix $\mathbf{\Sigma}_k$ the complexity and scalability of the model is worse compared to an model with univariate prior as in MOFA(+) [1, 2]. Due to the grid search approach on the lengthscales, we can cache for each grid point $p \in \{1, \ldots, P\}$ the corresponding covariance matrix $\mathbf{K}_k^{(C)}$ or its spectral decomposition that are required in each iteration. This leads to the following complexities for a model with a single group:

- initialization of grid points: $O(N^3P)$
- updates for $\mathbf{Z}$: $O(N^3K)$
- optimization of lengthscale and scale parameters: $O(PN^3K)$
- calculation of ELBO: $O(N^3K)$

A univariate prior on $\mathbf{Z}$ can reduce this to a quadratic complexity in N apart from the initialization step. The complexity in the number of features, views and factors stays linear as in the original model. As the scalability in the number of samples can be prohibitive for some applications, we below suggest an alternative more scalable approach using ideas from sparse Gaussian processes [12–14].

### 4 Scaling MEFISTO using sparse Gaussian processes

In order to avoid the cubic complexity in the number of samples, we can adopt the sparse Gaussian process framework [12–14] as previously suggested in the context of Gaussian Process Factor Analysis [15, 16]. For this, instead of considering all inputs we choose a set of M inducing points in order to capture the dependencies in the latent space. Here, we will choose a subset of the data points $S_m \subseteq (\{1, \ldots, N\})$ at locations $\tilde{\mathbf{C}} = \mathbf{C}_{S_m,:} \in \mathbb{R}^{M \times C}$ as inducing points and make inference on the remaining factor values conditional on the factor values at these points. For this, we denote the inducing points as $\mathbf{u}_k = z_k(\tilde{\mathbf{C}})$ and outline the resulting model below. We denote as above

$$(\mathbf{\Sigma}_k)_{nn'} = (1-\zeta_k) \exp\left(-\frac{||\mathbf{c}_n^g - \mathbf{c}_{n'}^{g'}||^2}{2\ell_k^2}\right) \kappa_k^G(g, g') + \zeta_k \delta_{n,n'} \tag{69}$$

or if the data has a Kronecker-structure

$$\mathbf{\Sigma}_k = (1-\zeta_k) \mathbf{K}^{(G)}_{\mathbf{x}_k,\sigma_k} \otimes \mathbf{K}^{(C)}_{\ell_k} + \zeta_k \mathbf{I}_{TG} \tag{70}$$

$$\text{with} \quad \mathbf{K}^{(C)}_{\ell_k} = \left(\exp\left(-\frac{||\mathbf{c}_t - \mathbf{c}_{t'}||^2}{2\ell_k^2}\right)\right)_{t,t'=1,\ldots,T} \tag{71}$$

$$\tilde{\mathbf{K}}^{(G)}_k = \sum_{r=1}^{R} \mathbf{x}_r^{(k)} \mathbf{x}_r^{(k)T} + \sigma_k^2 \mathbf{I}_G \tag{72}$$

As before, $\mathbf{\Sigma}_k$ depends on the Gaussian process hyperparameters. We further use $\mathbf{\Sigma}_{k,ZU}$ to denote the $N \times M$-submatrix obtained by keeping only the columns of $\mathbf{\Sigma}_k$ given by $S_m$ corresponding to the inducing points $\mathbf{U}$, analogously $\mathbf{\Sigma}_{k,UZ} = \mathbf{\Sigma}_{k,ZU}^T$ given by the rows of $\mathbf{\Sigma}_k$ in $S_m$ and we use $\mathbf{\Sigma}_{k,UU}$ to denote the $M \times M$-submatrix obtained by keeping only the columns and rows given by $S_m$. This leads to the following model:

$$p(y_{nd}^m | \mathbf{Z}, \mathbf{W}, \boldsymbol{\tau}) = \mathcal{N}\left(\sum_k w_{kd}^m z_{nk}, \tau_{md}^{-1}\right) \tag{73}$$

$$p(\mathbf{z}_{:,k} | \mathbf{u}_k) = \mathcal{N}(\mathbf{\Sigma}_{k,ZU} \mathbf{\Sigma}_{UU}^{-1} \mathbf{u}_k, \mathbf{\Sigma}_k - \mathbf{\Sigma}_{k,ZU} \mathbf{\Sigma}_{kUU}^{-1} \mathbf{\Sigma}_{k,UZ}) \tag{74}$$

$$p(\mathbf{u}_k) = \mathcal{N}(0, \mathbf{\Sigma}_{k,UU}) \tag{75}$$

and the remaining model components remain unchanged.

In particular, this means that we only need to model the covariance of factor values at selected input locations and make the inference for the remaining factor values conditional on these inducing points. Instead of the term $q(\mathbf{Z})$ in the variational distribution we now have a term $q(\mathbf{Z}, \mathbf{U})$, which we model as follows:

$$q(\mathbf{Z}, \mathbf{U}) = \prod_{k=1}^{K} q(\mathbf{z}_{:,k} | \mathbf{u}_k) q(\mathbf{u}_k) = \prod_{k=1}^{K} p(\mathbf{z}_{:,k} | \mathbf{u}_k) q(\mathbf{u}_k) \tag{76}$$

In particular, we take for the variational distribution of $\mathbf{Z}|\mathbf{U}$ the same density as for its prior.

#### 4.1 Updates

##### 4.1.1 Inducing points

To derive the updates for $\mathbf{u}_k$ we note the following dependency of the ELBO on $\mathbf{u}_k$:

$$\begin{aligned}
\mathcal{L}(q) &= \mathbb{E}_q[\log p(\mathbf{Y}, \mathbf{X})] - \mathbb{E}_q[\log q(\mathbf{X})] \\
&= \mathbb{E}_q[\log p(\mathbf{Y}, \mathbf{X})] - \mathbb{E}_q[\log q(\mathbf{z}_{:,k}, \mathbf{u}_k)] + const \\
&= \int q(\mathbf{u}_k) \int \log \frac{p(\mathbf{Y}, \mathbf{X})}{p(\mathbf{z}_{:,k} | \mathbf{u}_k) q(\mathbf{u}_k)} q(\mathbf{X}_{-\mathbf{u}_k}) d\mathbf{X}_{-\mathbf{u}_k} d\mathbf{u}_k + const \\
&= \int q(\mathbf{u}_k) \log \frac{\exp \mathbb{E}_{-\mathbf{u}_k} \log \frac{p(\mathbf{Y}, \mathbf{Z}, \mathbf{W}, \mathbf{U}, \boldsymbol{\alpha}, \boldsymbol{\theta}, \boldsymbol{\tau})}{p(\mathbf{z}_{:,k} | \mathbf{u}_k)}}{q(\mathbf{u}_k)} d\mathbf{u}_k + const
\end{aligned} \tag{77}$$

Therefore, for each factor $k$ we have

$$\log q^*(\mathbf{u}_k) = \mathbb{E}_{-\mathbf{u}_k} \log \frac{p(\mathbf{Y}, \mathbf{Z}, \mathbf{W}, \mathbf{U}, \boldsymbol{\alpha}, \boldsymbol{\theta}, \boldsymbol{\tau})}{p(\mathbf{z}_{:,k} | \mathbf{u}_k)}, \tag{78}$$

where the expectation is with respect to $q(\mathbf{X}_{-\mathbf{u}_k}) = q(\mathbf{W})q(\boldsymbol{\alpha})q(\boldsymbol{\tau})q()\boldsymbol{\theta})p(\mathbf{z}_{:,k}|\mathbf{u}_k)\prod_{l\neq k} q(\mathbf{z}_l, \mathbf{u}_l)$. This lead to the variational update being given by

$$q^*(\mathbf{u}_k) = \mathcal{N}(\boldsymbol{\nu}_k, \mathbf{B}_k), \tag{79}$$

where

$$\mathbf{B}_k = (\boldsymbol{\Sigma}_{k,UU}^{-1}\boldsymbol{\Sigma}_{ZU}^T \mathbf{S}\boldsymbol{\Sigma}_{k,ZU}\boldsymbol{\Sigma}_{k,UU}^{-1} + \boldsymbol{\Sigma}_{k,UU}^{-1})^{-1} \quad \text{with} \quad \mathbf{S} = \operatorname{diag}\left(\sum_{d,m}\langle w_{dk}^{(m)2}\rangle\langle\tau_d^{(m)}\rangle\right) \tag{80}$$

$$\boldsymbol{\nu}_k = \mathbf{B}_k\boldsymbol{\Sigma}_{k,UU}^{-1}\boldsymbol{\Sigma}_{k,ZU}^T\tilde{\mu} \quad \text{with} \quad \tilde{\mu} = \left(\sum_{m,d}\langle\tau_d^{(m)}\rangle\langle w_{dk}^{(m)}\rangle\left(y_{nd}^{(m)} - \sum_{l\neq k}\langle w_{dl}^{(m)}\rangle\langle z_{nl}\rangle\right)\right)_{n=1,\dots,N} \tag{81}$$

###### 4.1.2 Latent factors

By assumption on the variational distribution $q(\mathbf{Z}|\mathbf{U}) = p(\mathbf{Z}|\mathbf{U})$, we have for each factor $k$

$$q(\mathbf{z}_{:,k}|\mathbf{u}_k) = \mathcal{N}(\boldsymbol{\Sigma}_{k,ZU}\boldsymbol{\Sigma}_{k,UU}^{-1}\mathbf{u}_k, \boldsymbol{\Sigma}_k - \boldsymbol{\Sigma}_{k,ZU}\boldsymbol{\Sigma}_{UU}^{-1}\boldsymbol{\Sigma}_{k,UZ}). \tag{82}$$

As we only require the marginal distributions in the remaining updates, we calculate the mean and variance of the marginal distribution $q(z_{nk})$ given by

$$q(z_{nk}) = \mathcal{N}((\boldsymbol{\Sigma}_{k,ZU}\boldsymbol{\Sigma}_{k,UU}^{-1}\boldsymbol{\nu}_k)_n, (\boldsymbol{\Sigma} - \boldsymbol{\Sigma}_{ZU}\boldsymbol{\Sigma}_{k,UU}^{-1}\boldsymbol{\Sigma}_{k,UZ} + \boldsymbol{\Sigma}_{k,ZU}\boldsymbol{\Sigma}_{k,UU}^{-1}\mathbf{B}_k\boldsymbol{\Sigma}_{k,UU}^{-1}\boldsymbol{\Sigma}_{k,UZ})_{nn}) \tag{83}$$

###### 4.1.3 Remaining components (weight and noise terms)

The updates for terms in $w$, $\tau$, $\theta$, $\alpha$ remain unchanged. Note that they do not depend on $\mathbf{U}$ but on $\mathbf{Z}$ only. The required expectations of $\mathbf{Z}$ are given by from (83).

###### 4.1.4 Gaussian process hyperparameters

The optimization of the lengthscale and scale parameters of the sparse Gaussian process kernel is performed analogous to the full model by a grid search to maximize the ELBO term depending on $\ell_k$, $\zeta_k$ and (if multiple groups are included) $\mathbf{x}_k$ and $\sigma_k$, where here only the covariance matrix on the inducing points is required, thus reducing computational complexity:

$$\begin{aligned}\ell_k, \zeta_k, \mathbf{x}_k, \sigma_k &= \arg\max_{\ell_k,\zeta_k,\mathbf{x}_k,\sigma_k} \frac{1}{2}\log\det(\boldsymbol{\Sigma}_{k,UU}^{-1}) - \frac{1}{2}\langle\mathbf{u}_k^T\boldsymbol{\Sigma}_{k,UU}^{-1}\mathbf{u}_k\rangle + \frac{1}{2}\log\det(\mathbf{B}_k)\\ &= \arg\max_{\ell_k,\zeta_k,\mathbf{x}_k,\sigma_k} \frac{1}{2}\log\det(\boldsymbol{\Sigma}_{k,UU}^{-1}) - \frac{1}{2}\operatorname{Tr}(\boldsymbol{\Sigma}_{k,UU}^{-1}\mathbf{B}_k) - \frac{1}{2}\boldsymbol{\nu}_k^T\boldsymbol{\Sigma}_{k,UU}^{-1}\boldsymbol{\nu}_k + \frac{1}{2}\log\det(\mathbf{B}_k)\end{aligned} \tag{84}$$

Due to the grid search approach we again cache the $\mathbf{K}_{\ell_k}^{(C)}$ matrices or their spectral decomposition for each grid point, keeping an $N \times N$ matrix $\boldsymbol{\Sigma}_k$ containing both values of $\boldsymbol{\Sigma}_{k,UU}$ and $\boldsymbol{\Sigma}_{k,ZU}$ as sub-matrices. The inverse and its log-determinant values required throughout the updates need only be stored and calculated for the $M \times M$ matrix $\boldsymbol{\Sigma}_{k,UU}$, thus resulting in lower memory requirements and initialisation costs. As in the full inference, we decompose $\boldsymbol{\Sigma}_{k,UU}$ if possible as

$$\boldsymbol{\Sigma}_{k,UU} = (1-\zeta_k)\mathbf{K}_k^{(G)} \otimes \mathbf{K}_{k,UU}^{(C)} + \zeta_k\mathbf{I} \quad \text{with} \quad \mathbf{K}_{UU}^{(C)} = \left(\exp\left(-\frac{||\mathbf{c}_n - \mathbf{c}_{n'}||^2}{2\ell_k^2}\right)\right)_{n,n'\in S_m} \tag{85}$$

and use the spectral decomposition

$$\mathbf{K}_{k,UU}^{(C)} = \mathbf{V}_{k,UU}^{(C)}\mathbf{D}_{k,UU}^{(C)}\mathbf{V}_{k,UU}^{(C)T}. \tag{86}$$

This again allows a comparatively cheap evaluation of the objective function in the hyperparameters, as the matrices $\mathbf{V}^{(C)}$ and $\mathbf{D}^{(C)}$ can be precomputed at each grid point and the inverse and log determinant can be calculated in linear time complexity in the number of samples per group given the precomputed matrices as before. Note that this use of inducing points is mainly sensible in settings with a single group or a high numbers of samples per group, but not for data with many groups and a small number of time points per group.

#### 4.2 Evidence lower bound

In the evidence lower bound, the KL divergence term for $\mathbf{Z}|\mathbf{U}$ is zero, as the variational distribution was chosen identical to the prior, i.e. $\log q(\mathbf{z}_{:,k}|\mathbf{u}_k) = \log p(\mathbf{z}_{:,k}|\mathbf{u}_k)$. The KL divergence term for the inducing points can be found analogous to the term for the factors values in the non-sparse model with the sub-setted covariance matrix:

$$\langle \log p(\mathbf{u}_k) \rangle = \frac{1}{2} \log \det(\mathbf{\Sigma}_{k,UU}^{-1}) - \frac{1}{2} \langle \mathbf{u}_k^T \mathbf{\Sigma}_{k,UU}^{-1} \mathbf{u}_k \rangle - \frac{M}{2} \log(2\pi) \tag{87}$$

$$\langle \log q(\mathbf{u}_k) \rangle = -\frac{1}{2} \log \det(\mathbf{B}_k) - \frac{M}{2} - \frac{M}{2} \log(2\pi) \tag{88}$$

The remaining terms including the log likelihood term and KL divergence terms for the weight and noise parameters do not change. As noted above for the updates of noise and weight variables, these remaining ELBO terms again only depend on $\mathbf{Z}$ and not $\mathbf{U}$ and expectations for the factor values are taken from (83). As the new ELBO decomposes in $n$ except for the inducing points this sparse formulation would also open up application of stochastic inference.

#### 4.3 Choice of inducing points

The inducing points are chosen as a subset of the training inputs in a regular grid from the original locations where ties (e.g. from different sample groups) are shuffled randomly. The number of inducing points can be specified by the user, with a higher number resulting in slower but less approximative results. By default, we use a minimum of 50% of the input points as inducing points if this number is above 100, as for smaller values the full inference is still very fast. In principle one could also use pseudo-inputs at arbitrary locations and perform an optimization of the choice of inducing points (e.g. [14, 17]). This however comes with an increased computational burden.

### 5 Covariate alignment in the latent space using MEFISTO

If samples between different groups have no clear correspondence in their covariates such as developmental stages between species or disease progression time courses between patients, we need to align the samples between groups. To reduce the noise in the alignment of the sample, we implemented a procedure that enables simultaneously aligning and factorizing the data.

Given a one-dimensional covariate $c_n^g$ for $g = 1, \ldots, G$, $n = 1, \ldots N_g$ we interleave the updates in the model with an alignment step based on dynamic time warping [18] in the latent space with the possibility of partial alignment [19]. For this, the Euclidean distances between the expectations of the latent factor values $\langle z_{nk}^g \rangle$ under the current variational distribution are used as dissimilarity measure between two samples, i.e.

$$d(n_g, n'_{g'}) = ||\langle \mathbf{z}_n^g \rangle - \langle \mathbf{z}_{n'}^{g'} \rangle||_2 \tag{89}$$

to obtain a cross-distance matrix between two groups $g$ and $g'$. The dynamic time warping algorithm finds a warping curve $\omega = (\omega_g, \omega_{g'}) \in \{1, \ldots N_g\} \times \{1, \ldots N_{g'}\}$ that minimizes the distance between two groups by transforming the covariate axis of each group. This distance is given by

$$d_\omega(g, g') = \sum_{t=1}^{T} d(\omega_g(t), \omega_{g'}(t)) \frac{m_\omega(t)}{M_\omega}, \tag{90}$$

where $m_\omega(t)$ is a weighting function that defines the cost of an alignment and $M_\omega$ a normalization constant. Here, one group acts as the reference group and the other as a query group. To perform the alignment with more than two groups one can specify a reference group to the model, which is used as a reference to align all other groups. The warping function $\omega$ is constrained to provide a monotonic mapping from one group to the other group to maintain the order given by the covariate, i.e. $\omega_g(k+1) \geq \omega_g(k)\ \forall g, k$. Dynamic programming is used to find the optimal solution, the full procedure is implemented in the *dtw-python* package [18]. An illustration of the alignment is given in Figure 3.

By default, MEFISTO allows for partial matching with different end or beginning using an asymmetric step pattern that matches each element of the query group to exactly one element in the reference group. If the samples in each groups can be assumed to have the same begin and end point, this can be passed as an option to MEFISTO to perform a global alignment with complete matching using a symmetric step pattern, where a diagonal step has the same cost as two steps along each axis favouring diagonal steps in the alignment. Additional constrains could be implemented on the warping function apart from monotonicity, as for instance an admissible region given by a window around the diagonal.

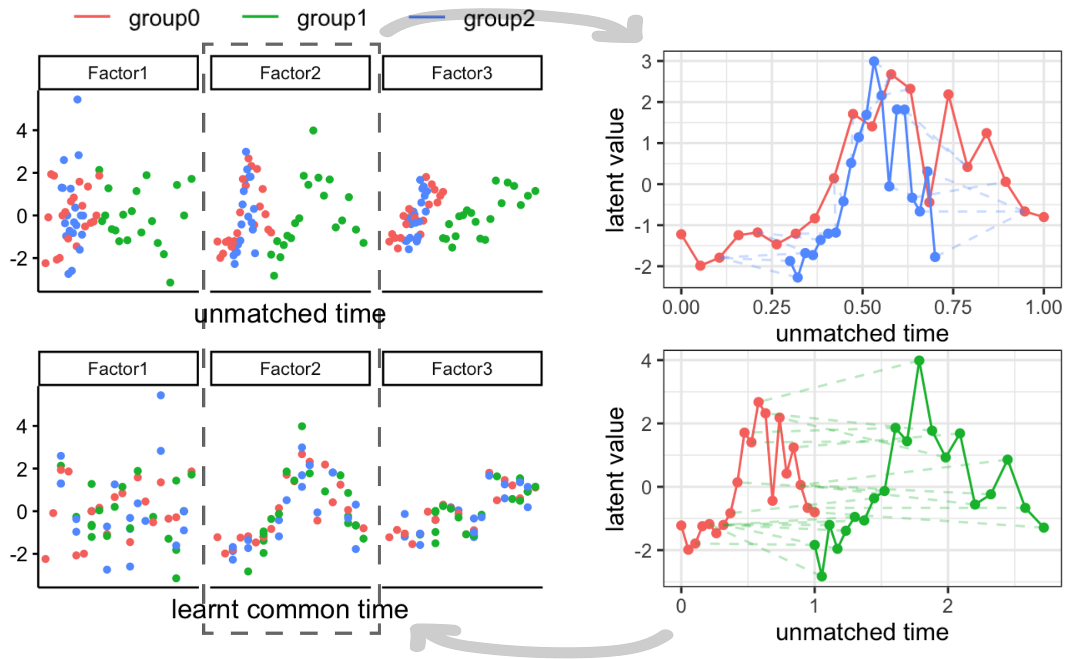

Figure 3: Example illustrating the alignment of a covariate (here time) between groups (indicated by different colours). The upper part of the left panel shows the factor values along time before alignment, the lower part after alignment. Times are matched to minimize the distance between groups across all factors by learning a monotonic warping function between groups. The right panel shows the alignment on Factor 2 in more detail for illustration purposes. Note, however, that the found alignment is based on all factors jointly.

### 6 Down-stream analyses

Based on the weights of the model, similar down-stream analyses can be conducted as in MOFA2 [2], including feature set enrichment analysis and inspection of the top weights per factor, in order to uncover the molecular drivers of variation that underlie a latent factor. Similarly clustering and outlier identification can be performed in the latent space. The functional view on the latent factors however opens up additional analyses such as separation of smooth from non-smooth factors, inspection of group relationships per factor using the inferred group kernel matrices or interpolation or extrapolation of factors.

#### 6.1 Smoothness and sharedness scores per factor

**Separation of smooth and non-smooth patterns** The hyperparameters of the model directly give insights into the smoothness of a factor. A smoothness score per factor is based on $s_k = 1 - \zeta_k$. This value ranges from 0 indicating a non-smooth factor to 1 indicating a very smooth factor.

**Inspection of group relationships and sharedness scores per factor** With multiple groups, the group-group kernel given by $\mathbf{K}_k^G$ can be used to cluster the groups or identify outliers on the level of groups for each latent factor $k$. An overall sharedness score per factor is calculated based on the mean absolute distance to the identity covariance (no-sharedness) in the off-diagonal elements. A value of 1 indicates sharedness, a value of 0 no sharedness between groups for the given factor.

##### 6.2 Interpolation & extrapolation

In some settings, we might obtain new covariate values and would like to predict the value that a sample would take in the latent space without having any data available for this sample. For example, in developmental studies we might be interested in non-observed time points during development or in longitudinal clinical studies in the extrapolation of the disease course for future time points. While such predictions into unseen regions need to be always taken with care, the Gaussian process framework enables to make such prediction and at the same time provides measures of uncertainty associated with the predictions. Given a new value of the covariate $\mathbf{c}^* \in \mathbb{R}^C$ we obtain the posterior of the corresponding latent factor value $\mathbf{z}^* \in \mathbb{R}^K$ as

$$p(z_k^*|\mathbf{Y}) = \mathcal{N}\left(\kappa(\mathbf{c}^*, \mathbf{C})\mathbf{\Sigma}_k^{-1}\boldsymbol{\nu}_k, \kappa(\mathbf{c}^*, \mathbf{c}^*) + \zeta_k - \kappa(\mathbf{c}^*, \mathbf{C})\mathbf{\Sigma}_k^{-1}\kappa(\mathbf{C}, \mathbf{c}^*) + \kappa(\mathbf{c}^*, \mathbf{C})\mathbf{\Sigma}_k^{-1}\mathbf{A}_k\mathbf{\Sigma}_k^{-1}\kappa(\mathbf{C}, \mathbf{c}^*)\right).$$

The resulting predictions and uncertainties can be visualized as illustrated in Figure 4. Given the latent prediction we can further impute actual values of the measured features from the generative model underlying MEFISTO.

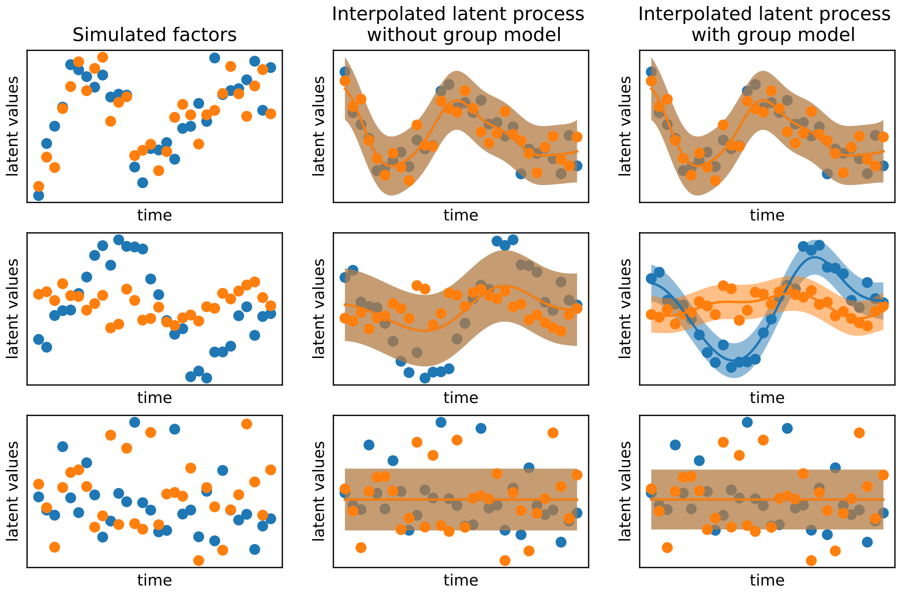

Figure 4: Illustration of interpolation: The left column shows the simulated factors for observed time points, the middle column the continuous interpolation without respecting the group structure, the right column shows the interpolation if the sharedness is learnt by the model and group structure are taken into account for interpolation. The line denotes the predictive mean of the process, the shaded region the 95% confidence interval.

### 7 Comparison to related approaches

MEFISTO is based upon (sparse) factor analysis models that have previously been used in genomics, both for data sets comprising a single feature set [20–23] as well as for multi-modal data sets that consist of several feature sets and sample groups [1, 2]. These methods however do not model continuous structures among samples which naturally occur for example in temporal or spatial data. Here, we account for such structures by employing Gaussian process priors in the latent space. This use of Gaussian processes is related to previous approaches in neurobiology, geostatistics or image analysis [15, 16, 24, 25], where a Gaussian process prior is used to model a latent space constructed by a linear or non-linear mapping from the observed data. Existing models in these fields however mostly consider a single view and group [16, 24] or multiple views that share the exact same weights (and features) [15].

| Method | inference | sample structure | sparsity of weights | scalable functional model | multiple views | noise model | temporal alignment | mapping |
|---|---|---|---|---|---|---|---|---|
| GPFA [16] | VI | continuous | No | Yes | No | Gaussian | No | linear |
| GPFA [24] | EM | continuous | No | No | No | Gaussian | No | linear |
| svGPFA [15] | VI | continuous | No | Yes | only for identical features | Gaussian, Poisson process | Yes | linear |
| MOFA+ [2] | VI | groups | feature- and view-wise | n/a | Yes | Gaussian, Poisson, Bernoulli | n/a | linear |
| MOFA [1] | VI | None | feature- and view-wise | n/a | Yes | Gaussian, Poisson, Bernoulli | n/a | linear |
| GPPVAE [25] | VI | continuous + groups | No | Yes | No | Gaussian | No | non-linear (VAE) |
| Dependent PMF [26] | MCMC | continuous | No | No | only for identical features | Gaussian | No | linear |
| timeOmics [27] | MLE and subsequent PCA | continuous | feature- and view-wise | Yes | Yes | Gaussian | No | linear |
| GPVAE [28] | VI | continuous | feature-wise | Yes | No | Gaussian | No | non-linear (VAE) |
| DLGFA [29] | VI | continuous | view-wise | Yes | Yes | Gaussian | No | non-linear (RNN) |
| CTF [30] | alternating least squares | None | No | n/a | No | None | No | linear |
| MEFISTO | VI | continuous + groups | feature- and view-wise | Yes | Yes | Gaussian, Poisson, Bernoulli | Yes | linear |

Table 1: Table for comparison of MEFISTO to related approaches. EM stands for Expectation-Maximization,VI for variational inference, VAE for variational auto-encoder, RNN for recurrent neural network.

Furthermore, these models do not incorporate sparsity constrains on weights or views and mostly do not account for multiple groups of samples. A more detailed comparison is provided in Table 1. In addition, some methods have considered factor models for temporal or spatial data using two-step approaches, e.g. interpreting temporal relationships post-hoc once the factors have been learned [30] or first smoothing the observations along time and then applying factor models [27]. These approaches can however not make use of the functional nature of the factor, which is for example required for interpolation, extrapolation or separation of smooth from non-smooth factors. In addition, they do not provide a model of group heterogeneity or an alignment procedure and thus can suffer from high variation between groups.

### References

1. Argelaguet, R. *et al.* Multi-Omics factor analysis - a framework for unsupervised integration of multi-omic data sets. *Molecular Systems Biology* (2018).
2. Argelaguet, R. *et al.* MOFA+: A statistical framework for comprehensive integration of multi-modal single-cell data. *Genome Biology* **21,** 1–17 (2020).
3. Virtanen, S., Klami, A., Khan, S. & Kaski, S. *Bayesian group factor analysis* in *Artificial Intelligence and Statistics* (2012), 1269–1277.
4. Klami, A., Virtanen, S., Leppäaho, E. & Kaski, S. Group factor analysis. *IEEE transactions on neural networks and learning systems* **26,** 2136–2147 (2015).
5. Rasmussen, C. E. & Williams, C. K. I. *Gaussian processes for machine learning* (MIT press Cambridge, 2006).
6. Rakitsch, B., Lippert, C., Borgwardt, K. & Stegle, O. It is all in the noise: Efficient multi-task Gaussian process inference with structured residuals. *Advances in Neural Information Processing Systems,* 1–9 (2013).
7. Bishop, C. M. Pattern recognition. *Machine Learning* **128,** 1–58 (2006).
8. Murphy, K. P. *Machine learning: a probabilistic perspective* (2012).
9. Blei, D. M., Kucukelbir, A. & McAuliffe, J. D. Variational Inference: A Review for Statisticians. *Journal of the American Statistical Association* **112,** 859–877. arXiv: `1601.00670` (2017).
10. Svensson, V., Teichmann, S. A. & Stegle, O. SpatialDE: identification of spatially variable genes. *Nature Methods* **15,** 343–346 (2018).
11. Arnol, D., Schapiro, D., Bodenmiller, B., Saez-Rodriguez, J. & Stegle, O. Modeling Cell-Cell Interactions from Spatial Molecular Data with Spatial Variance Component Analysis. *Cell Reports* **29,** 202–211.e6 (2019).
12. Hensman, J., Fusi, N. & Lawrence, N. D. Gaussian Processes for Big Data. *Proceedings of the Twenty-Ninth Conference on Uncertainty in Artificial Intelligence, Corvallis, OR: AUAI Press,* 282–290 (2013).
13. Matthews, D. G. *et al.* GPflow: A Gaussian process library using TensorFlow. *Journal of Machine Learning Research* **18,** 1–6 (2017).
14. Titsias, M. K. Variational Learning of Inducing Variables in Sparse Gaussian Processes. *Proceedings of Machine Learning Research* **5,** 567–574 (2009).
15. Duncker, L. & Sahani, M. Temporal alignment and latent Gaussian process factor inference in population spike trains. *Advances in Neural Information Processing Systems,* 10445–10455 (2018).
16. Luttinen, J. & Ilin, A. Variational Gaussian-process factor analysis for modeling spatio-temporal data. *Advances in Neural Information Processing Systems 22 - Proceedings of the 2009 Conference,* 1177–1185 (2009).
17. Bauer, M., Van Der Wilk, M. & Rasmussen, C. E. Understanding probabilistic sparse Gaussian Process approximations. *Advances in Neural Information Processing Systems,* 1533–1541. arXiv: `1606.04820` (2016).
18. Giorgino, T. Computing and visualizing dynamic time warping alignments in R: The dtw package. *Journal of Statistical Software* **31,** 1–24 (2009).
19. Tormene, P., Giorgino, T., Quaglini, S. & Stefanelli, M. Matching incomplete time series with dynamic time warping: an algorithm and an application to post-stroke rehabilitation. *Artificial Intelligence in Medicine* (2009).
20. Witten, D. M., Tibshirani, R. & Hastie, T. A penalized matrix decomposition, with applications to sparse principal components and canonical correlation analysis. *Biostatistics* **10,** 515–534 (2009).
21. Gehring, J. S., Fischer, B., Lawrence, M. & Huber, W. SomaticSignatures: Inferring mutational signatures from single-nucleotide variants. *Bioinformatics* **31,** 3673–3675 (2015).
22. Stegle, O., Parts, L., Piipari, M., Winn, J. & Durbin, R. Using probabilistic estimation of expression residuals (PEER) to obtain increased power and interpretability of gene expression analyses. *Nature Protocols* **7,** 500–507 (2012).

23. Alexandrov, L. B., Nik-Zainal, S., Wedge, D. C., Campbell, P. J. & Stratton, M. R. Deciphering Signatures of Mutational Processes Operative in Human Cancer. *Cell Reports* **3,** 246–259 (2013).

24. Yu, B. M. *et al.* Gaussian-process factor analysis for low-dimensional single-trial analysis of neural population activity. *Journal of Neurophysiology* **102,** 614–635 (2009).

25. Casale, F. P., Dalca, A. V., Saglietti, L., Listgarten, J. & Fusi, N. Gaussian Process Prior Variational Autoencoders. *Advances in Neural Information Processing Systems* (2018).

26. Adams, R. P., Dahl, G. E. & Murray, I. Incorporating side information in probabilistic matrix factorization with Gaussian processes. *Proceedings of the 26th Conference on Uncertainty in Artificial Intelligence, UAI 2010,* 1–9. arXiv: 1003.4944 (2010).

27. Bodein, A., Chapleur, O., Droit, A. & Lê Cao, K. A. A Generic Multivariate Framework for the Integration of Microbiome Longitudinal Studies With Other Data Types. *Frontiers in Genetics* **10,** 1–18 (2019).

28. Fortuin, V., Baranchuk, D., Rätsch, G. & Mandt, S. *GP-VAE: Deep Probabilistic Time Series Imputation* in (eds Chiappa, S. & Calandra, R.) **108** (PMLR, Online, 2019), 1651–1661. arXiv: 1907.04155.

29. Qiu, L., Chinchilli, V. M. & Lin, L. Deep Latent Variable Model for Longitudinal Group Factor Analysis. arXiv: 2005.05210 (2020).

30. Martino, C. *et al.* Context-aware dimensionality reduction deconvolutes gut microbial community dynamics. *Nature Biotechnology* (2020).
